# Structural insights into perilipin 3 membrane association in response to diacylglycerol accumulation

**DOI:** 10.1101/2022.11.17.516819

**Authors:** Yong Mi Choi, Dalila Ajjaji, Kaelin D. Fleming, Peter P. Borbat, Meredith L. Jenkins, Brandon E. Moeller, Shaveen Fernando, Surita R. Bhatia, Jack H. Freed, John E. Burke, Abdou Rachid Thiam, Michael V. Airola

## Abstract

Lipid droplets (LDs) are dynamic organelles that contain an oil core mainly composed of triglycerides (TAG) that is surrounded by a phospholipid monolayer and LD-associated proteins called perilipins (PLINs). During LD biogenesis, perilipin 3 (PLIN3) is recruited to nascent LDs as they emerge from the endoplasmic reticulum. Here, we analyzed how lipid composition affects PLIN3 recruitment to membrane bilayers and LDs, and the structural changes that occur upon membrane binding. We found the TAG precursors phosphatidic acid and diacylglycerol (DAG) recruit PLIN3 to membrane bilayers and define an expanded Perilipin-ADRP-Tip47 (PAT) domain that preferentially binds DAG enriched membranes. Membrane binding induces a disorder/order transition of alpha helices within the PAT domain and 11-mer repeats, with intramolecular distance measurements consistent with the expanded PAT domain adopting a folded but dynamic structure upon membrane binding. In cells, PLIN3 is recruited to DAG enriched ER membranes, and this requires both the PAT domain and 11-mer repeats. This provides molecular details of PLIN3 recruitment to nascent LDs and identifies a function of the PAT domain of PLIN3 in DAG binding.

## INTRODUCTION

Lipid droplets (LDs) act as energy reservoirs in cells. They contain a neutral lipid core of mainly triacylglycerols (TAGs) and cholesterol esters with a phospholipid monolayer that surrounds the neutral lipid core. This creates a membrane environment for LDs that is distinct from membrane bilayers and recruits several LD-associated proteins [1–3]. In addition to a major role in energy storage, LDs are also important cellular hubs that traffic proteins and lipids between organelles, regulate ER stress, and contribute to viral infections [4–6].

LDs are formed in the ER where neutral lipids are synthesized [7, 8]. Mechanistically, LD formation involves several steps [9] including accumulation of neutral lipids in the outer ER membrane leaflet, neutral lipid nucleation aided by the seipin complex and associated factors (e.g. LDAF1) [10], formation of nascent LDs that bud from the ER [11, 12] and LD growth and maturation [13, 14]. Several lines of evidence support the idea that the phospholipid composition of the ER membrane is locally edited to promote LD assembly or recruit specific proteins important for LD formation such as seipin and ORP proteins [7, 15–17]. Although there is still uncertainty regarding the phospholipid composition at the initial stages of LD formation, evidence from yeast suggest diacylglycerol (DAG), the direct precursor of TAG, is enriched in discrete ER subdomains where LD biogenesis is initiated [15, 18]. DAG can also promote TAG nucleation and impact the architecture of LDs on the ER [15, 19].

Perilipins (PLINs) are the major class of proteins that coat the surface of LDs [20–26]. There are five PLINs in humans that bind LDs at various stages of their initiation and maturation [21]. For example, PLIN1 is the major PLIN that binds to mature LDs in adipocytes, while PLIN2 and PLIN5 reside on mature LDs in liver and muscle cells, respectively [27]. In contrast, PLIN3 displays near ubiquitous expression and binds to early LDs as they bud from the ER but is later displaced by other PLINs as LDs grow and mature [10, 28]. PLIN3 is stable and not degraded in the absence of LDs. This allows PLIN3 to translocate from the cytoplasm to sites of early LD formation, where PLIN3 is well established to act as a marker for the biogenesis of early LDs across species [10, 28, 29].

PLINs share a conserved protein domain architecture composed of an N-terminal PAT domain, followed by variable stretches of 11-mer repeats and a C-terminal 4-helix bundle [30, 31]. The 11- mer repeats form amphipathic helixes that are sufficient to recruit PLINs to LDs [30–32], while the 4-helix bundle in some PLINs can also bind LDs [30, 33]. The binding of amphipathic helices, such as those found in the 11-mer repeats, to membrane interfaces is greatly influenced by the level or presence of phospholipid packing defects [34, 35]. The degree of packing is determined by the type of lipid present (as indicated in the triangle in **Fig. 1A**). When it comes to the oil-water interface of LDs, the recruitment of amphipathic helices is greater when the level of phospholipid packing is lower [35, 36], i.e. when there are more packing defects. Although the PAT domain is the most conserved domain among PLINs, its function is not clear, as it has not been demonstrated to bind to membranes or LDs.

**Figure 1.**
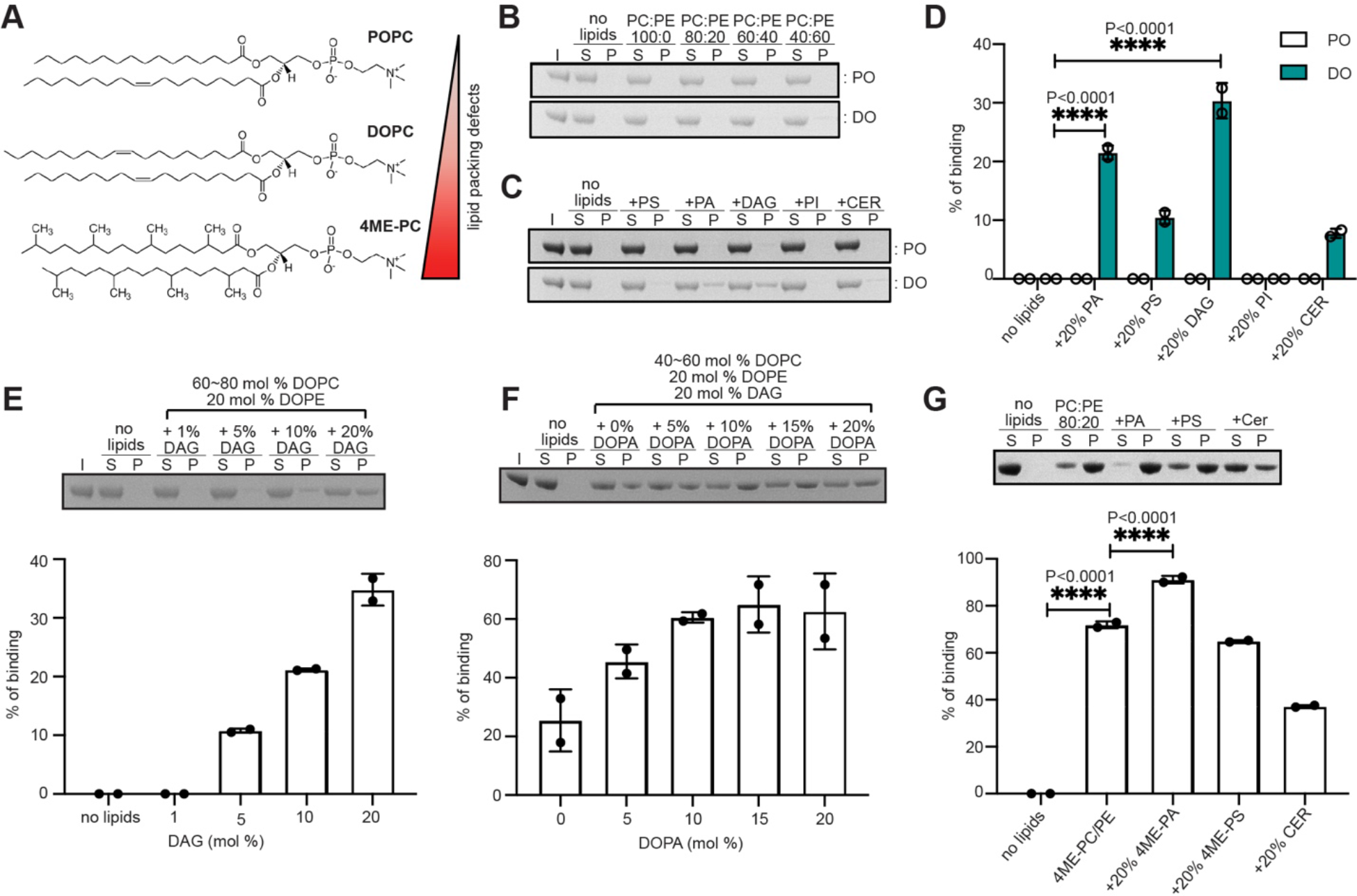
Triglyceride precursors DAG and PA recruit PLIN3 to membrane bilayers **A)** Structures of representative phospholipids used in this study. **B)** SDS-PAGE analysis of human PLIN3 liposome sedimentation assay on the either palmitoyl- oleoyl (PO)-liposomes or di-oleoyl (DO)-liposomes in different ratio of PC to PE. Lane S (supernatant) and P (pellet) represent unbound and bound human PLIN3 to liposomes. **C)** SDS-PAGE analysis of human PLIN3 recruitment to liposomes enriched with 20 mol% of the additional lipids PS, PA, DAG, PI and ceramide. **D)** Quantification of PLIN3 recruitment to liposomes in different lipid compositions (n=2). Statistical analysis was performed using two-way ANOVA (****, p<0.0001). **E)** SDS-PAGE analysis of human PLIN3 recruitment to DO-liposomes using different amounts of DAG. DOPE concentration was held constant at 20mol% and DOPC was adjusted between 60 and 80mol% to maintain total phospholipid concentration. Bar graph shows addition of DAG increases PLIN3 binding. **F)** SDS-PAGE analysis of human PLIN3 recruitment to DO-liposomes enriched with different amount of DOPA. The amounts of DOPE and DAG were kept constant at 20mol%. DOPC concentration was adjusted between 40 and 60mol% to maintain total phospholipid concentration. Bar graph shows addition of DOPA further increases PLIN3 binding. **G)** SDS-PAGE analysis of human PLIN3 recruitment to 4ME liposomes enriched with 20 mol% of the additional lipids 4ME-PA, 4ME-PS, ceramide. 4ME-PC and 4ME-PE concentrations ranged between 60 ∼80mol% and 20mol%, respectively. Statistical analysis for quantification of liposome binding was performed using ordinary one-way ANOVA with Tukey’s multiple comparison test (n=2, ****,p<0.0001)

Here, we examine the mechanism of PLIN3 recruitment to membrane bilayers and LDs using a combination of *in vitro* and cell culture assays, and analyze the structural changes induced by membrane binding using hydrogen-deuterium exchange mass spectrometry (HDX-MS) and pulsed-dipolar electron spin resonance spectroscopy (PD-ESR). We found that human PLIN3 is recruited to membrane bilayers enriched in the TAG precursors phosphatidic acid (PA) and DAG. By delineating the roles of the PAT domain and 11-mer repeats, we define an expanded PAT domain that is sufficient for PLIN3 to bind DAG enriched membranes, while the 11-mer repeats are sufficient to bind LD monolayers. We confirm that DAG enrichment can drive PLIN3 recruitment to ER membranes in cells, and this requires both the PAT domain and 11-mer repeats. Structurally, the PAT domain and 11-mer repeats form inducible alpha helices to drive membrane association and the PAT domain forms a tertiary structure upon membrane binding. Taken together, this study provides molecular insight into how PLIN3 is recruited to early LDs and reveals a novel function for the PAT domain of PLIN3 in DAG binding.

## RESULTS

### The Triglyceride Precursors DAG and PA Recruit PLIN3 to Membrane Bilayers

To systematically define how lipid composition affects PLIN3 recruitment to membrane bilayers, we purified recombinant human PLIN3 from *Escherichia coli* and used liposome co-sedimentation assays to monitor membrane binding. Liposomes were prepared using multiple freeze/thaw cycles and characterized by dynamic light scattering (DLS) **(Fig. S1A)**. Liposomes were made from phospholipids with different acyl-chain combinations comprised of either palmitoyl-oleoyl (PO) or di-oleoyl (DO) phospholipids **(Fig. 1A)**. PO phospholipids have one unsaturated acyl chain and are representative of the typical lipid composition in ER membranes. DO phospholipids contain two unsaturated acyl chains, have increased membrane packing defects [34, 37–39] and are enriched in ER membranes after oleate supplementation, which stimulates LD formation [15, 40–42]. Initial liposome co-sedimentation experiments varied the ratio of neutral phospholipids phosphatidylcholine (PC) and phosphatidylethanolamine (PE), as PC and PE represent the major lipids on both the cytoplasmic face of the ER and surface of LDs, PE increases both PLIN2 binding to liposomes [43] and PLIN3 insertion into mixed lipid monolayers at phospholipid-oil interfaces [44] and the ratio of PC to PE has previously been shown to regulate protein distribution on LDs [43, 45]. Under all PC-to-PE ratios tested, PLIN3 did not bind liposomes comprised solely of PC and PE **(Fig. 1B)**.

We next asked whether addition of other lipids that are synthesized in the ER could recruit PLIN3 to membranes by generating PC/PE liposomes containing 20mol% of either phosphatidic acid (PA), phosphatidylserine (PS), diacylglycerol (DAG), phosphatidylinositol (PI) or C18:1 ceramide. Regardless of lipid composition, PLIN3 did not bind to any PO-based liposomes **(Fig. 1C, 1D)**. However, PLIN3 was recruited to DO-based liposomes enriched in DAG or PA [37, 38, 46] **(Fig. 1C, 1D)**. Addition of DOPS and ceramide resulted in a minor increase in PLIN3 binding in DO-based liposomes, while the addition of PI had no effect **(Fig. 1C, 1D)**.

PLIN3 recruitment to DO-based liposomes depended on the surface concentration of DAG, with 5mol% DAG able to induce ∼10% binding, and 20mol% DAG inducing 35% binding **(Fig. 1E)**. The addition of DOPA further increased PLIN3 binding in DAG-containing liposomes, which suggested a synergistic effect of PA and DAG on PLIN3 membrane recruitment **(Fig. 1F)**. Increasing the total liposome concentration caused the majority of PLIN3 to bind membranes, but 5% of PLIN3 remained in the soluble fraction **(Fig. S2A)**. DLS characterization indicated DAG enriched liposomes had two populations based on size **(Fig. S1C, S1D)**, which is likely due to the fusogenic properties of DAG [46]. The addition of DOPA or PLIN3 protein decreased or eliminated the population of the larger (>400 nm) liposomes **(Fig. S1C, S1D)**. We concluded that PLIN3 binds liposomes containing DAG and/or PA, which are notably the membrane-lipid precursors for triglyceride synthesis, and PLIN3 can also remodel membranes, consistent with previous studies [29].

### PLIN3 Binds to Liposomes with Membrane Packing Defects

LDs have increased membrane packing defects in comparison to membrane bilayers [47]. Previously, PLIN4 has been shown to have increased binding to liposomes composed of methyl branched diphytanoyl (4ME) phospholipids that create shallow lipid-packing defects that can be accessed by hydrophobic insertion of peripheral membrane proteins, such as PLINs [31, 48]. The packing defects induced by 4ME phospholipids are greater than DO phospholipids because of the wider space between lipids [48]. In comparison to DO and PO liposomes, 4ME phospholipids significantly increased PLIN3 liposome association with ∼70% binding observed with a mixture of 4ME-PC and 4ME-PE **(Fig. 1G)**. Consistent with our previous observations, 4ME-PA further increased PLIN3 binding, while 4ME-PS and C18:1 ceramide decreased binding **(Fig. 1G)**. The effect of DAG was unable to be assessed as addition of DAG to 4ME-PC/PE disrupted proper liposome formation. Taken together, this suggests PLIN3 membrane binding is also dependent on lipid packing defects, and PA can further enhance PLIN3 membrane binding.

### PLIN3 Binding to Artificial Lipid Droplets

As the major function of PLIN3 is to bind to emerging LDs as they bud from the ER, we next assessed the ability of recombinant PLIN3 to bind artificial lipid droplets (ALDs) *in vitro* using a flotation assay [10, 28, 29, 49]. ALDs were generated with a triolein neutral lipid core surrounded by a phospholipid monolayer of DOPC and DOPE in 1:2.5 molar ratio of phospholipids to TAG. ALDs with 80mol% DOPC and 20mol% DOPE showed 50% binding of PLIN3 **(Fig. S2B)**. The addition of the anionic phospholipids PI4P decreased PLIN3 binding to ALDs. In contrast, PA, PS and PI slightly decreased binding, ceramide did not affect binding, and DAG slightly increased PLIN3 binding. However, none of these effects were statistically significant. The modest impact of these PA, PS, PI, DAG, and ceramide, which all can induce negative curvature or charge in membrane bilayers, may be due to the basal occurrence of significant phospholipid packing voids on the ALD surface as compared with bilayers [36, 47]; while the strongly charged PI4P would instead mask these and diminish binding [47].

### The PAT Domain and 11-mer Repeats are Disordered in the Absence of Membranes

We next sought to examine how the structure of PLIN3 changes upon membrane binding, first focusing on the structure of PLIN3 in the absence of membranes. For these experiments, we used hydrogen deuterium exchange mass spectrometry (HDX-MS), which measures the exchange of amide hydrogens with deuterated solvent. This method acts as a readout for protein conformational dynamics with regions that form secondary structures undergoing slower deuterium exchange than disordered regions, which lack intramolecular hydrogen bonds and secondary structure [50, 51]. A brief pulse of deuterated solvent is useful for identifying regions within a protein that lack structure compared to ordered regions [52, 53].

We determined the absolute exchange of PLIN3 after a 3sec pulse of deuterium incorporation at 0°C (equivalent to ∼0.3sec at 20°C) using a fully deuterated control. After the deuterium pulse, the first 200 residues of PLIN3 comprising the PAT domain and 11-mer repeats were fully deuterated, indicating these regions are completely disordered in the absence of membranes **(Fig. 2A)**. The 4-helix bundle and *α*/*β* domain were largely ordered with comparatively low rates of deuterium incorporation **(Fig. 2A)**, which is consistent with prior structural studies of PLIN3 [54, 55].

**Figure 2.**
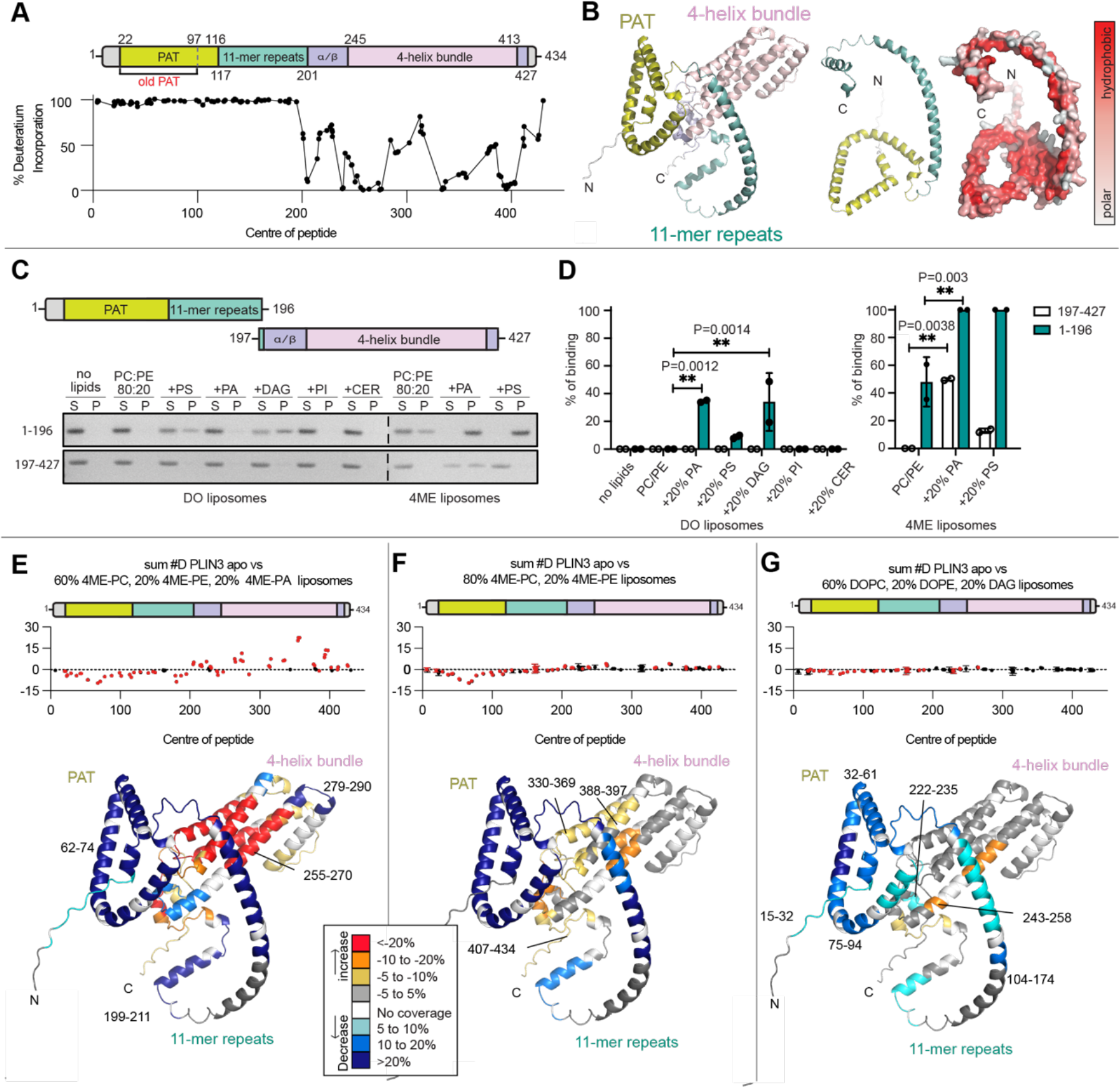
HDX-MS analysis of PLIN3 membrane binding **A)** Absolute percentage of deuterium incorporation after 3sec deuterium exposure of PLIN3 at 1°C in the absence of liposomes. Each point represents a single peptide, with them being graphed on the x-axis according to their central residue. A domain architecture of human PLIN3 was drawn to match the scale of x-axis of HDX-MS. N-terminal PAT domain (mustard) and 11- mer repeats (cyan) have no detectable secondary structure in the absence of liposomes. **B)** A predicted 3D structure of human PLIN3 (UniProt: O60664) by AlphaFold (Left). The surface of PAT domain and 11-mer repeats were shown according to the hydrophobicity (Right). **C)** SDS-PAGE and **D)** quantitative analysis of PAT/11-mer repeats and 4-helix bundle liposome binding. DO and 4ME-liposomes were prepared in the same way as Figure 1C and 1D. Statistical analysis for quantification of liposome binding was performed using two-way ANOVA with Tukey’s multiple comparison test (n=2). **E-G)** Quantitative analysis of deuterium exchange differences of human PLIN3 in the presence of liposomes. The sum of the difference in the # of incorporated deuterons is shown for the absence and the presence of liposomes over all timepoints. The N-terminus composed of PAT/11-mer repeats was significantly protected from deuterium exchange (defined as > 5% change in exchange, > 0.4 Da mass difference in exchange, a *p*-value < 0.01 using a two-tailed Student’s *t*-test). Each point represents an individual peptide, with those colored in red having a significant difference, with error bars showing standard deviation (n=3). Liposomes were generated with **(E)** 60mol% 4ME-PC, 20mol% 4ME-PE and 20mol% 4ME-PA, **(F)** 80mol% 4ME- PC, 20mol% 4ME-PE or **(G)** 60mol% DOPC, 20mol% DOPE and 20mol% DAG. A map of deuterium exchange rate according to all peptides throughout the entire PLIN3 was generated based on the AlphaFold predicted PLIN3 structure and color coded according to the legend. The full set of peptides is shown in the source data.

### The PAT Domain and 11-mer Repeats are the Major Drivers of PLIN3 Membrane Association

We next sought to determine any conformational changes that occur during membrane binding using HDX-MS. Having established optimal conditions for PLIN3 membrane binding, we measured the deuterium exchange rate over various time points (3, 30, 300, and 3000sec) in the absence or presence of 4ME liposomes composed of 60mol% 4ME-PC, 20mol% 4ME-PE, and 20mol% 4ME-PA.

In the presence of membranes, striking differences in deuterium exchange were observed throughout the PLIN3 sequence **(Fig. 2E, Table S1, source data)**. The most notable differences were large decreases in deuterium exchange in the PAT domain and 11-mer repeat regions. In comparison to previous results that demonstrated the 11-mer repeats are sufficient for the lipid droplet association [30, 31], this suggests that the PAT domain and 11-mer repeats are both major contributors to membrane binding. Consistently, we found that a purified PAT/11-mer repeats fragment displayed similar membrane recruitment as full length PLIN3 to DO liposomes enriched in DAG and PA, and 4ME liposomes **(Fig. 2C, 2D)**.

The HDX-MS results suggested that membrane binding induces the formation of a secondary structure in the PAT domain and 11-mer repeats. This is consistent with both AlphaFold and RoseTTAFold models that predict the PAT domain and 11-mer repeats form amphipathic alpha helices, with the PAT domain adopting a triangular globular structure and the 11-mer repeats forming a series of extended alpha helices **(Fig. 2B, 3B)**. Based on these results we concluded that the PAT domain and 11-mer repeats of PLIN3 are intrinsically disordered in the absence of membranes, with inducible amphipathic alpha helices being stabilized upon membrane binding.

**Figure 3.**
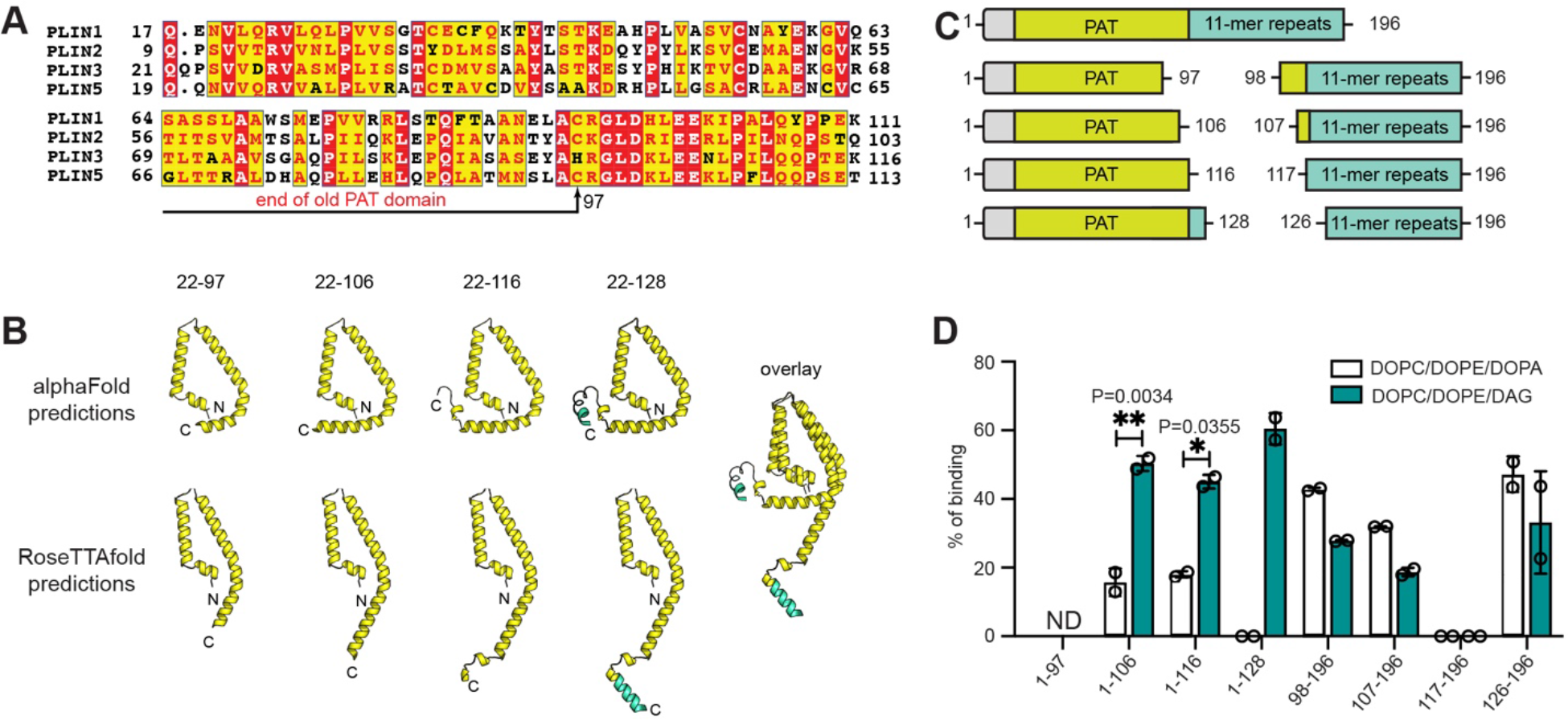
An expanded PAT domain binds DAG enriched membranes **A)** Multiple sequence alignment of the N-terminus of human PLIN 1 (UniProt: O60240), PLIN2 UniProt: Q99541), PLIN3 (UniProt: O60664), and PLIN5 (UniProt: Q00G26). The previously suggested PAT domain (residues 1-97) is indicated with black arrow. **B)** Putative PAT domain triangular structures predicted by both AlphaFold and RoseTTAFold. **C)** Schematic of PAT domain constructs and the counterpart 11-mer repeats constructs. **D)** Quantitative analysis of liposome recruitment for various PAT domain and 11-mer repeats constructs. Statistical analysis for quantification of liposome binding was performed using two- way ANOVA with Tukey’s multiple comparison test (n=2).

Notably, many of the peptides with the PAT domain and 11-mer repeats displayed a bimodal distribution of deuterium exchange upon membrane binding **(Fig. S3A, S3B)**. This differs from a canonical distribution where the degree of deuteriation is gaussian in individual peptides. The observed bimodal distributions are consistent with either EX1 exchange [56], or there being two distinct protein conformations in the sample. Depending on the membrane residency of PLIN3, bimodal distributions can be consistent with observations of the free and membrane bound state. Given our previous observation that the PAT and 11-mer repeats are intrinsically disordered in the absence of membranes, the simplest interpretation is that the PAT domain and 11-mer repeats cycle between two states: 1) a membrane bound alpha helical state on membranes that is strongly protected from H-D exchange and 2) an intrinsically disordered solution state that rapidly exchanges with deuterated solvent upon dissociation from membranes and leads to rapid H-D exchange. Therefore, the half-life of bimodal exchange can act as a surrogate for membrane residency time for disordered regions of the protein.

The protection from deuterium exchange and the non-gaussian bimodal distribution for most peptides in the PAT domain persisted over a longer time course (∼300sec) in comparison to peptides within the 11-mer repeat regions (∼30sec) **(Fig. S4A)**. This trend of longer HDX protection correlated with the degree of sequence conservation among PLINs **(Fig. 3A)**. Taken together this supports a role for the PAT domain in membrane binding and suggests a similar role for the highly conserved PAT domain in other PLINs.

In contrast to the N-terminal PAT domain and 11-mer repeats, the C-terminal 4-helix bundle displayed a large increase in deuterium exchange **(Fig. 2E, S4A)**. The most apparent increase in deuterium exchange was in the middle of the 4-helix bundle. We hypothesized this was due to the 4-helix bundle unfolding upon membrane binding, with this possibly contributing to membrane recruitment. To test this, we examined the ability of a purified PLIN3 4-helix bundle fragment (residues 197-427) to bind membranes. Consistent with our hypotheses, the isolated 4-helix bundle bound to the 4ME-PC/PE/PA liposomes that were used in the HDX-MS experiment **(Fig. 2C, 2D)**. However, the 4-helix bundle did not bind to 4ME-liposomes lacking PA, or to any DO- based liposomes even when PA and DAG were present **(Fig. 2C, 2D)**. We concluded that the 4- helix bundle of PLIN3 can unfold and bind membranes, but only under very specific condition such as the presence of accumulated PA on the membrane having packing defects induced by 4ME-PC/PE. This conclusion is supported by the PA accumulation at the nascent LD formation stie where lipid packing defects occur [17].

To understand how lipid composition influences the structure of PLIN3, we again applied HDX- MS, using variable liposome compositions of 4ME-PC/PE and DOPC/PE/DAG **(Fig. 2F, 2G)**. In line with of previous HDX-MS and liposome sedimentation experiments, we observed protection from H-D exchange in the PAT domain and 11-mer repeats using 4ME-PC/PE and DOPC/PE/DAG liposomes, with the magnitude and duration of H-D exchange protection in these regions correlating with the observed fraction of PLIN3 bound in liposome sedimentation assays **(Fig. S4B, 1D and 1G)**.

In contrast, with the 4ME-PC/PE and DOPC/PE/DAG lipid compositions, the 4-helix bundle did not display large increases in deuterium exchange indicating that 4-helix bundle is not involved in membrane binding in the absence of PA. Taken together, these results confirm that the PAT domain and 11-mer repeats are the major lipid interacting regions of PLIN3, and that the 4-helix bundle forms a stable tertiary structure that in general does not contribute to membrane association, unless PA is present in membrane areas with shallow lipid packing defects. This condition is particularly fulfilled at nascent LD formation sites.

### An expanded PAT domain binds DAG enriched membranes

Our results suggested a role for the PAT domain in PLIN3 membrane association. However, several studies [30, 31] have previously established the 11-mer repeats of PLINs are sufficient for membrane binding, and to our knowledge, the PAT domain alone has not been demonstrated to bind membranes. Complicating matters, the precise boundaries of the PAT domain have been difficult to establish. Previous studies have suggested the PAT domain consists of the N-terminal 97 residues in PLIN3 [30], which are highly conserved with other PLINs **(Fig. 3A)**. However, high sequence conservation between PLINs continues beyond residue 97 and ends at residue 115 in PLIN3 **(Fig. 3A)**. This suggests the PAT domain may be larger than previously expected. Consistent with the sequence conservation, both Alphafold [57] and RoseTTA fold [58] predict a triangular alpha helical tertiary structure for the PAT domain that encompasses residues 22-116 **(Fig. 3B)**.

To investigate the PAT domain boundaries and membrane interaction capabilities, we purified several C-terminal extended PAT domain constructs and the corresponding 11-mer repeats counterparts **(Fig. 3C)** and assessed the ability of these fragments to bind DO liposomes containing either 20mol% PA or 20mol% DAG. As an important note, we were unable to purify the previously suggested PAT domain construct (residues 1-97), as this construct was not stably expressed in *Escherichia coli*. This is consistent with the poor expression of the PAT domain of PLIN1 in yeast [30].

In general, fragments encompassing the PAT domain bound strongly to liposomes in the presence of DAG, with modest liposome binding in the presence of PA **(Fig. 3D)**. In contrast, the 11-mer repeats bound to liposomes containing either PA or DAG **(Fig. 3D)**. Taken together, the data suggests that the PAT region does form a domain that spans residues 22-116 in PLIN3. Notably, this expanded PAT domain is capable of membrane binding and displays a preference for binding DAG enriched membranes, while the 11-mer repeats of PLIN3 do not.

### Conformational Rearrangements of the PAT Domain upon Membrane Binding

We first sought to confirm the effects of liposome binding on secondary structure of PLIN3 by circular dichroism (CD) **(Fig. S5)**. Liposomes were prepared with 4ME-PC/PE/PA, which recruited all the PLIN3 constructs (full length, PAT, 11-mer repeats, PAT/11-mer repeats, 4-helix bundle) in the previous liposome co-sedimentation experiments. For full length PLIN3, the presence of liposomes increased overall helicity as observed by an increased negative peak around 222nm in the CD spectra. In comparison, the CD spectra of the 4-helix bundle was largely unaffected by the presence of liposomes and was consistent with a stable alpha helical structure. In contrast, the PAT/11-mer repeats adopted a mostly random coil structure in solution with a negative peak around 200nm, and a shift to alpha helices in the presence of liposomes as indicated by a large negative peak at 222nm. Liposomes induced similar changes in the CD spectra for both the PAT domain and 11-mer repeats alone. Taken together, this confirms that the increase in helicity observed in full length PLIN3 by membranes was due to the PAT/11-mer repeats undergoing a disorder/alpha helical transition. This is in line with our HDX-MS results and prior studies of PLIN2 and PLIN3 fragments [55, 59].

Next, to investigate if membranes induced conformational changes in the PAT domain, we applied the pulsed-dipolar spectroscopy (PDS) technique of double electron-electron resonance (DEER) [60, 61] to full length PLIN3 in solution and bound to liposomes. PDS is a collection of several, based on recording electron spin-echo (ESE), pulse ESR techniques [62–66] routinely applied to characterize protein conformations by providing accurate constraints in the distance range of ∼10- 90Å. The amplitude of detected ESE depends on dipolar coupling between the spins of unpaired electrons in nitroxide spin labels covalently bound to engineered cysteine residues. Stepping out pulse separation in the sequence produces the time domain ESE envelope, from which the distances could be reconstructed. [62–65].

Two sets of residues (25 and 96; 37 and 114) were selected for spin labeling, which were respectively 71 and 77 residues apart in the primary sequence but predicted to be in close proximity by AlphaFold and RoseTTAFold **(Fig. 4A-4C)**. Comparison of the DEER decoherence (decay) times indicated that the PAT domain is less structured and/or intrinsically disordered in solution versus when bound to liposomes, which is consistent with our HDX-MS and CD analysis **(Fig. 4D)**. One can estimate, based on ∼20% extent of decay in solution compared to liposomes that the decoherence time is of the order of 20μs, which corresponds to nearly a 100Å separation expected for a random polypeptide chain with stiffness [67]. In contrast, with liposomes the decoherence times are fast enough to decay to background well within the evolution time of 1.2 μs used in the liposome DEER measurements. The decays do not show oscillations, as is frequently observed for spin pairs with narrow distance distributions [68–71]. As a control to rule out lateral aggregation [69], we attached a single spin-label to residue 37C and did not observe a dipolar evolution (DEER shape) that could indicate a pair with a shorter than ∼80Å separation based on only slightly concave shape of the signal **(Fig. 4E)**. The DEER data thus suggest that conspicuous dipolar signals in the doubly labeled proteins result from intramolecular interactions, rather than being caused by intermolecular interactions on membranes.

**Figure 4.**
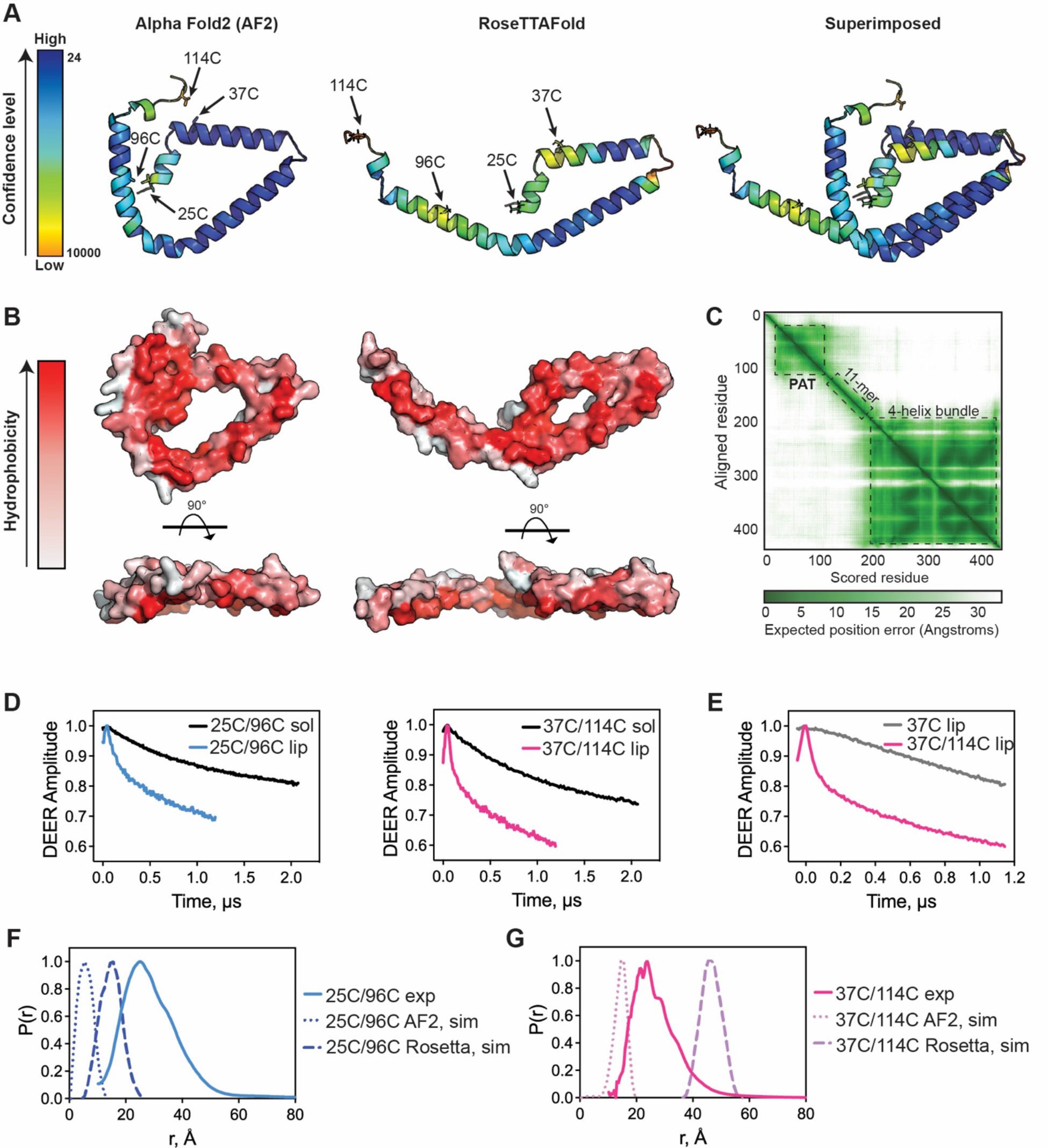
Membrane-induced conformational rearrangements are consistent with a PAT domain tertiary structure **A)** Cartoon representation of the PAT domain structure predicted by AlphaFold2 and RoseTTAFold with relative confidence levels shown. The positions of the spin-labeled residues are indicated with arrows. **B)** The hydrophobic surface of predicted PAT domain are depicted. The hydrophobic face of the helices is facing towards the reader. **C)** The PAE (Predicted Aligned Error) value of AlphaFold2 for full length PLIN3 was plotted by ChimeraX and shown as an interactive 2D plot (bottom, right panel). **D)** Time domain signals from DEER spectroscopy of double-labeled 25C/96C (left) and 37C/114C (right) in solution (sol) and on liposomes (lip). **E)** Time domain signals from DEER spectroscopy of double-labeled 37C/114C and single- labeled 37C proteins on liposomes. **F, G)** Distance distributions from DEER spectroscopy of **(E)** 25C/96C and **(F)** 37C/114C determined experimentally (exp) and from MtsslWizard simulations using the AlphaFold2 (AF2, sim) and RoseTTAFold (Rosetta, sim) predicted structures.

Distance distributions could be obtained in the presence of liposomes. The results showed semi- broad distance spreads of ∼20-40Å for the 25C/96C pair and ∼20-35Å for the 37C/114C pair **(Fig. 4F, 4G)**. These distances are comparable but do not exactly match the simulated distances using the AlphaFold and RoseTTAFold structure predictions that have only modest confidence levels **(Fig. 4A)** [57, 58]. For comparison, an extended alpha helix would result in distances of ∼100Å for these two sites. We also checked the spectral shape by recording field-swept echo with pulse separation of 250ns and did not notice any conspicuous broadening that could indicate a shorter range of distances (<15Å). We do however see from continuous wave (CW) ESR of 37C/114C **(Fig. S6)** that there might be a sizeable fraction of spins in the 15-20Å range whose contribution to the distance distribution will be significantly attenuated, since DEER has low sensitivity to distances in this range. The conformations with this distance range correlate well with AlphaFold predictions. The spread of the P(r) to longer distances could originate from the mobility of the C-terminal helix where residue 114C is located **(Fig. 4C)**. Taken together, we concluded that the PAT domain does form a folded domain when bound to membranes and this domain is likely mobile with a structure similar but not identical to the AlphaFold and RoseTTAFold predictions.

### DAG recruits PLIN3 to membrane bilayers and droplets in vitro

To verify a role for the PAT domain in DAG binding, we conducted an independent set of experiments to test if purified PLIN3 with GFP fused to the N-terminus bound DAG enriched membranes using droplet embedded vesicles (DEVs) [7, 35, 45, 72]. DEVs have emerged as a powerful in vitro system to model emerging LDs that bud from membrane bilayers. A major advantage of DEVs is the ability to not only monitor membrane association, but to also examine the protein distribution and preference for droplet monolayers versus membrane bilayers.

DEVs were generated by adding the neutral lipids TAG or DAG to giant unilamellar vesicles (GUVs) of DOPC that were marked with fluorescent phospholipids **(Fig. 5A)**. As observed previously [45, 73], full length PLIN3 was strongly recruited to the monolayer surface of triolein droplets with almost no bilayer signal **(Fig. 5B, 5D)**. Addition of diolein (DO-DAG) generated small droplet buds, and full length PLIN3 was recruited equally to the droplet and bilayer surface **(Fig. 5C, 5D)**. Residues 1-204 of PLIN3 that contained both the PAT domain and 11-mer repeats behaved nearly identical to full length PLIN3 in both TAG and DAG containing DEVs **(Fig. 5B, 5C and 5E)**.

**Figure 5.**
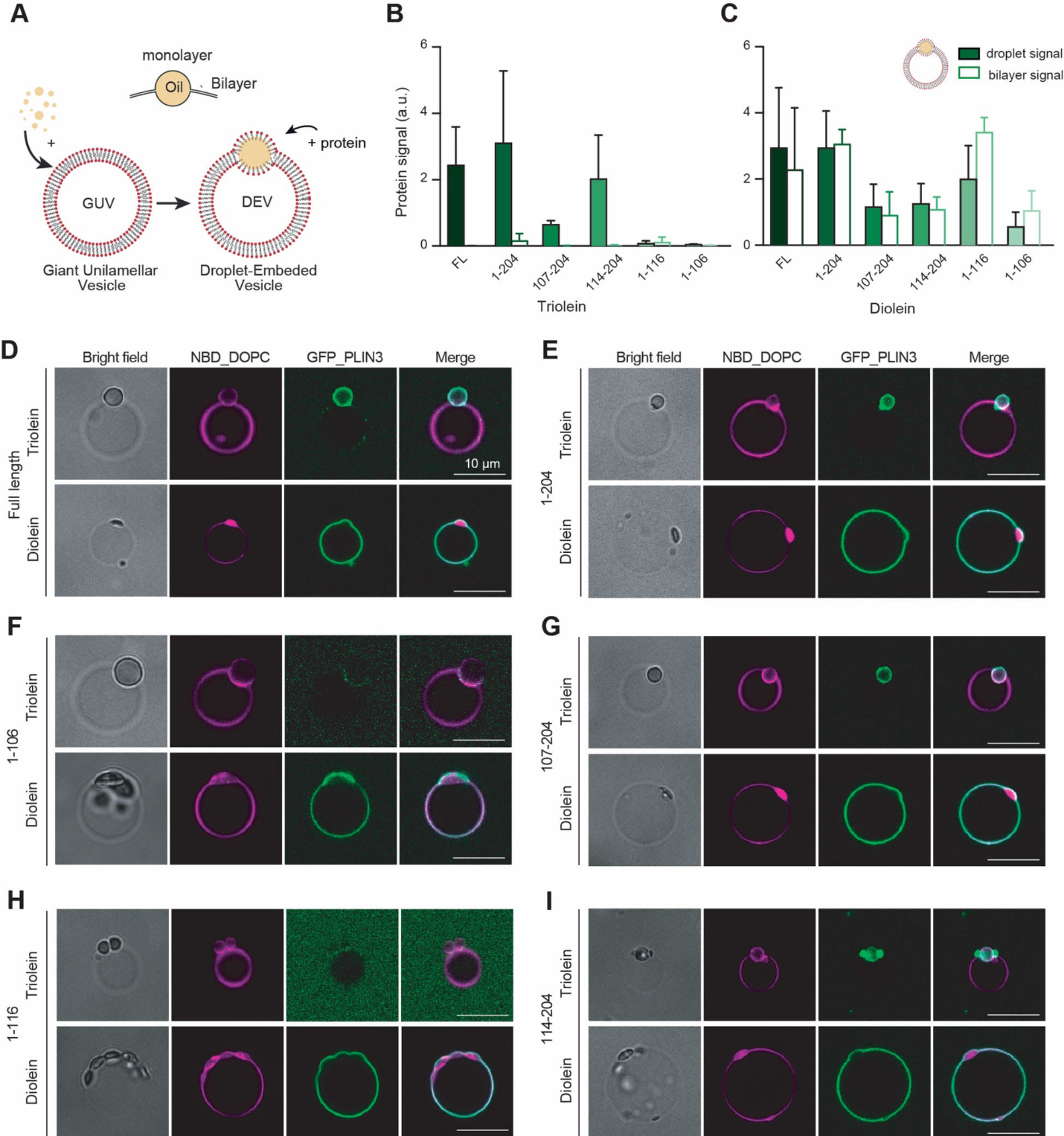
DAG recruits PLIN3 to membrane bilayers and LD droplets in vitro **A)** A diagram of generation of DEVs by mixing oils and giant unilamellar vesicles (GUVs). **B, C)** Quantification of the recruitment of GFP tagged PLIN3 constructs to droplet embedded vesicles (DEVs) containing (B) triolein or (C) Diolein. Statistical analysis was done with a Mann– Whitney non-parametric test. P<0.01 for all proteins. Three different experiments were performed and 10 to 15 DEVs quantified for each experiment. **D-I)** Fluorescent microscopic images of N-terminal GFP tagged full length and PLIN3 constructs (green) comprised of PAT domain and 11-mer repeats to (DEVs) (magenta) that are generated with fluorescent labeled phospholipids and oils. F) and H) Recruitment of PAT domain constructs. Almost no recruitment of PAT domain constructs in the presence of Triolein. G) and I) Recruitment of 11-mer repeats constructs.

Next, we tested fragments of the PAT domain and 11-mer repeats using both triolein and diolein DEVs. The 11-mer repeats preferentially bound the droplet surface of triolein DEVs with the magnitude of signal dependent on the length of the 11-mer repeats **(Fig. 5B, 5G and 5I)**. In contrast, PAT domain fragments did not bind to the droplet or bilayer surface of triolein DEVs **(Fig. 5B, 5F and 5H)**. The PAT domain was recruited to diolein DEVs, and the largest fragment (residues 1-116) bound to both the droplet and bilayer surface with similar magnitudes to full length PLIN3 **(Fig. 5C, 5F and 5H)**. The 11-mer repeats also bound to both the droplet and bilayer surface of diolein DEVs, but to a lesser extent **(Fig. 5C, 5G and 5I)**. These results are consistent with our previous hypothesis that the PAT domain is larger than previously expected and that this functional PAT domain binds DAG enriched membranes, while the 11-mer repeats can also bind DAG enriched membranes but display a preference for TAG containing droplets over membrane bilayers.

### DAG accumulation is sufficient to recruit PLIN3 to the ER in cells

DAG has previously been proposed to recruit PLIN3 to the ER in cells by blocking its hydrolysis or acylation or by the exogenous addition of DAG [74]. This is consistent with the current model for LD formation, where DAG accumulates at the site of TAG nucleation on the ER membrane [18, 19, 75], the local high concentration of neutral lipids promotes LD nucleation through seipin [15], and cytoplasmic PLIN3 marks these sites [10, 49]. We sought to test whether DAG accumulation was sufficient for PLIN3 recruitment using an independent system and also assess what fragments of PLIN3 were necessary and sufficient for ER recruitment.

To best visualize PLIN3 recruitment to ER membranes, we generated intracellular giant ER vesicles (GERVs) by submitting cells to hypotonic medium [72, 76, 77] **(Fig. 6A)**. After exchange to hypotonic medium, cells were pretreated with DMSO or the DGAT inhibitors (DGATi), followed by oleic acid to induce TAG or DAG synthesis [77] **(Fig. 6A)**. Confocal microscopy was used to visualize cells prior and after exchange to hypotonic medium, and after oleic acid treatment. Imaging was done in the following minutes after the treatments.

**Figure 6.**
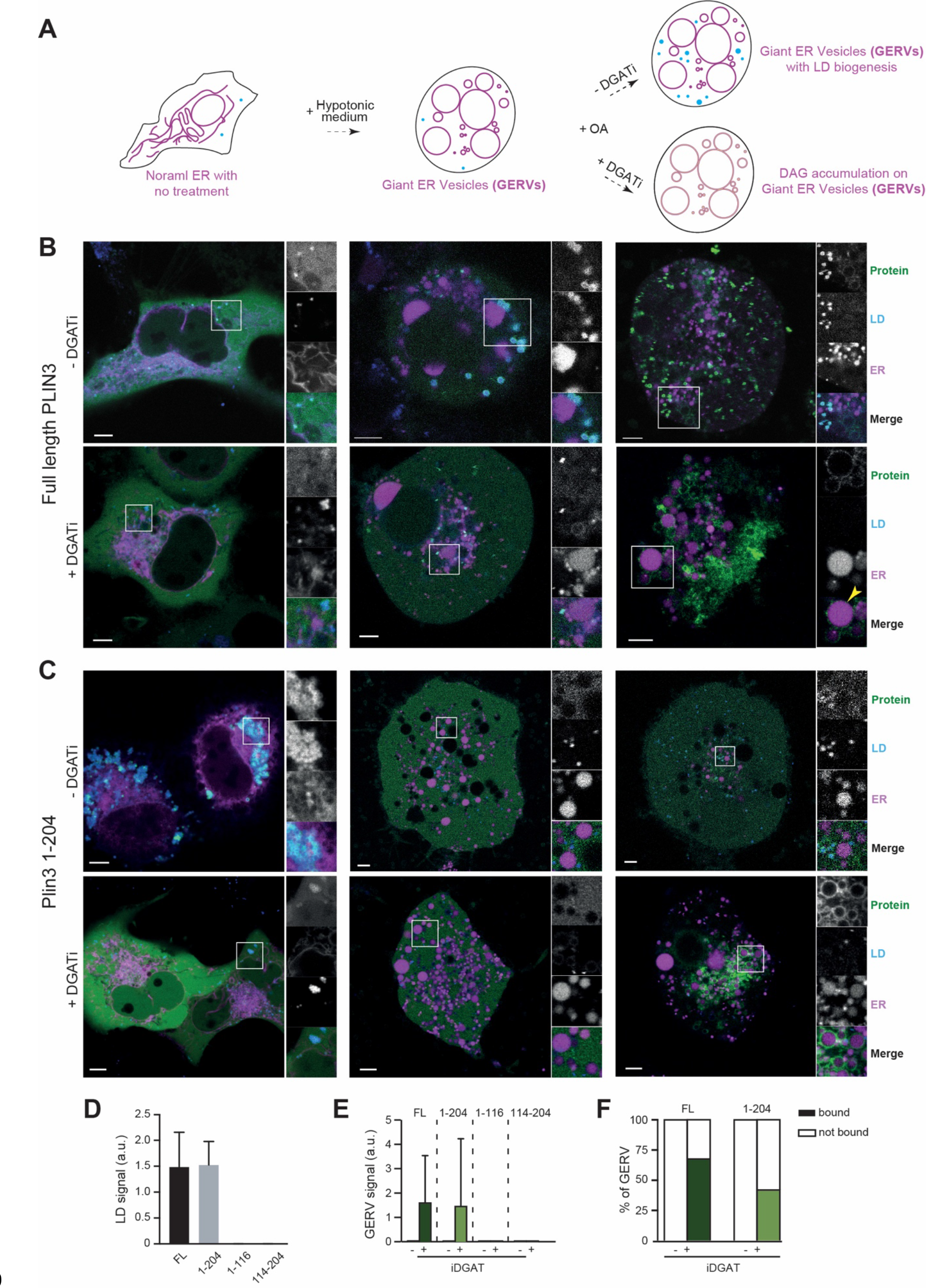
Both PAT domain and 11-mer repeats are necessary for PLIN3 recruitment to DAG enriched ER membranes in Cos7 cells **A)** A diagram of treatment of Giant ER Vesicle formation and treatment of oleic acid and DGATi to induce DAG accumulation on the ER in Cos7 cells. **B, C)** Subcellular localization of GFP tagged full length PLIN3 and PAT/11-mer repeats (1-204) were visualized in green in Cos7 cells under fluorescent microscope ZEISS LSM800 Airyscan. ER was visualized with ER specific marker RFP-KDEL in magenta. Lipid droplets were labeled with LipidTox Deepred to stain neutral lipids in cyan. After treated with hypotonic medium, cells were supplemented with oleate in the presence or absence of DGAT1/2 inhibitors. Each experiment was performed more than 3 times. **D)** Quantification of the amount of various PLIN3 constructs on LDs **E)** Quantification of the amount of various PLIN3 constructs on GERV after oleate treatment in the presence and the absence of DGAT1/2 inhibitors. **F)** % of GERV that are covered with full length PLIN3 or PAT/11-mer repeats construct (1-204) after oleate treatment in the presence and the absence of DGAT1/2 inhibitors.

As expected, the majority of subcellular localization of GFP-tagged PLIN3 was cytoplasmic in both normal and hypotonic media without addition of oleic acid **(Fig. 6B, upper panels)**, and co- localized with LDs **(Fig. 6D)**. Under conditions of DGAT inhibition, PLIN3 co-localized to the outer periphery of GERVs **(Fig. 6B, yellow arrow, 6E and 6F)**, which suggests DO-DAG accumulation is sufficient to recruit PLIN3 to ER membrane bilayers.

In line with our previous results, a construct containing both the PAT domain and 11-mer repeats showed a similar subcellular localization as full length PLIN3 under all conditions **(Fig. 6C-6F)**. This suggests the PAT domain and 11-mer repeats are sufficient for PLIN3 recruitment to both DAG enriched membrane bilayers and TAG containing LDs, which might also contain DAG whose concentration could increase the protein binding level. Interestingly, constructs containing only the PAT domain (residues 1-116) or only the 11-mer repeats (residues 114-204), which bound to DEVs in a DAG-dependent manner, remained cytoplasmic under all conditions **(Fig. 6D, 6E, S7)**. Taken together, we concluded that PLIN3 is capable of binding both DAG enriched ER membranes and early LDs, and that the PAT domain and 11-mer repeats are both necessary and synergize to perform these functions in cells.

## DISCUSSION

Here we find that PLIN3 not only binds LDs, but also membrane bilayers enriched in DAG and PA. These results are in line with other studies that revealed PLIN3 binding to DAG [74, 78, 79]. In our hands, DAG binding was observed with multiple in vitro systems (e.g. liposomes, DEVs) and DAG was also sufficient for PLIN3 recruitment to the ER in cells. Cellular recruitment to membranes by DAG is consistent with a seminal study that used inhibitors to block DAG hydrolysis or acylation and exogenous DAG to promote ER recruitment of PLIN3 [74]. Independent experiments in yeast have also demonstrated that membrane-anchored PLIN3 is sufficient to bind to DAG enriched subdomains in the ER [78, 79]. Here we show that oleate addition in combination with DGAT inhibition is sufficient for the subcellular redistribution of PLIN3 to the ER, which mimics normal the LD biogenesis pathway.

There are several mechanistic implications of PLIN3 recruitment to DAG enriched membranes. First, this implies that PLIN3 is recruited to sites of LD formation, not only through TAG generation, but also at the initial stages when DAG begins to accumulate. This raises a likely possibility that PLIN3 may play an active role in the early stages of LD formation by stabilizing DAG enriched regions on ER [78], and define sites of LD formation, before or in concomitant with seipin. Once PLIN3 binds to accumulated DAG, it might coat the curved surface of a growing DAG/TAG lens and regulate LD budding, in conjunction with seipin.

PA is another TAG precursor that is present at sites where LD originate [17] and can bind to seipin [16]. In addition to DAG, it was found to have a significant impact on the recruitment of PLIN3 to membranes in vitro. Thus, it may increase the translocation of PLIN3 to early LD formation sites. These two TAG precursors appear to provide specificity for the association of PLIN3 with membranes. From a curvature standpoint, DAG has a more negative curvature compared to PA and PE, whereas PE has a more negative curvature than PA [80]. This suggests that membrane curvature cannot solely account for the major role of PA in PLIN3 membrane binding specificity. However, from a surface charge perspective, PA and PE together may act synergistically by increasing charge of PA on the membrane [46, 81, 82]. In this scenario, PLIN3 recruitment to LD nucleation sites could be enhanced by specific recognition of PA, potentially through the 4-helix bundle. Therefore, PLIN3 membrane association may not only be determined by membrane packing defects, but could also involve selective physical interactions between PLIN3 and DAG or PA. This idea is also supported by a previous study that found that the LD binding properties of PLINs are sensitive to the polar residue composition of their amphipathic helices [83].

In this study, we attempted to clarify the function and boundaries of the PAT domain. Our HDX-MS results using full length PLIN3 clearly implicate both the PAT domain and 11-mer repeats in membrane binding. In addition, we were able to define a functional PAT domain that encompasses all of the conserved residues within PLINs and is longer than previously suggested. This expanded PLIN3 PAT domain is sufficient to bind DAG enriched membranes, but not LDs. In contrast, the 11-mer repeats display some affinity for DAG enriched membranes and are necessary to bind LD monolayers. Our overall conclusion is that the PAT domain and 11-mer repeats serve synergistic functions, as the individual regions are necessary for both LD and DAG recruitment in cells.

The PAT domain is predicted to adopt a triangular tertiary structure by both AlphaFold and RoseTTAFold. The DEER distance measurements and CW ESR data are not identical to these predictions, but do indicate that when bound to membranes the PAT domain adopts a folded domain. This conclusion is supported by our HDX-MS results that found the peptides within the PAT domain display longer protection times from H-D exchange, which could be due to either a tertiary structure more resistant to unfolding or a longer membrane residency time. We note that the membrane bound PAT domain structure is likely dynamic and additional distance measurements at distinct sites are needed to verify the accuracy of the predicted triangular structures.

To our knowledge, our finding that the PAT domain of PLIN3 is sufficient to bind DAG represents the first functional role for the PAT domain in any PLIN. This suggests that the highly conserved PAT domain in other PLINs may serve similar or related functions. For example, could the PAT domain be a general sensor for DAG in membrane bilayers? Or could the PAT domain of other PLINs potentially bind to other neutral lipids (e.g. CEs, TAGs, retinol esters)? Given the previously observed differing neutral lipid preference of PLINs [20] and the involvement of the PAT domain in lipid/membrane binding, this now raises the questions if this is due to the PAT domain preference, the 11-mer repeat preference, or the synergistic action of the combined PAT/11-mer repeat units.

Lastly, our results are consistent with previous studies that found the PLIN3 4-helix bundle does not bind LDs, but other PLINs 4-helix bundles can bind LDs [33]. While the domain boundaries of the 4-helix bundles seem reasonable given structure of mouse PLIN3 [54], the variable membrane binding affinities we observed for the PAT domain and 11-mer repeats does raise the question if stability of 4-helix bundle, and consequently the ability to unfold and bind membranes or LDs in other PLINs may depend on the domain boundaries of specific constructs. Thus, we suggest it is reasonable to revisit the ability of the 4-helix bundles from other PLINs using constructs that can be stably purified in vitro to rule out potential artifacts from the use of unstable 4-helix bundles.

## MATERIALS AND METHODS

### Protein expression and purification

The genes encoding PLIN3 constructs were codon-optimized for expression in *Escherichia coli* and cloned into pTHT, which is a derivative of pET28 that contains a TEV cleavable N-terminal 6x His tag. For GFP tagged PLIN3 constructs, monomeric superfolder GFP (msfGFP) was inserted between the 6x His tag and the N-terminus of PLIN3. PLIN3 plasmids were transformed into BL21(DE3) RIPL cells and protein expression was induced with 1mM isopropyl β-D-1- thiogalactopyranoside (IPTG) at 37°C for 3h. Cells was harvested by centrifugation at 3,320 x g for 20mins and stored at -80°C until use. Cell pellets were resuspended with buffer A containing 500mM NaCl, 20 mM Tris-HCl pH 7.5, 5% glycerol, 5mM 2-mercaptoethanol (BME) and lysed by sonication. Cell lysates were centrifuged at 81,770 x g for 1h at 4°C and the resulting supernatant was incubated with 5mL of Ni-NTA resin for 2h at 4°C prior to loading onto a gravity column. The Ni-NTA resin was washed with buffer B containing 500mM NaCl, 20mM Tris-HCl pH 7.5, 5% glycerol, 5mM BME and 60mM imidazole. Human PLIN3 proteins were eluted with buffer C containing 500mM NaCl, 20mM Tris-HCl pH 7.5, 5% glycerol, 5mM BME and 300mM imidazole. Eluted protein was applied to a HiLoad 26/600 Superdex 200 pg column (GE life sciences) equilibrated with 150mM NaCl, 20mM Tris-HCl pH 7.5, 5% glycerol, and 5mM BME. The average yield of full length PLIN3, PAT/11-mer repeats and 4-helix bundle throughout expression and purification was between 5-10mg from a 1L culture of *Escherichia coli.* The yield of PAT or 11- mer repeats constructs or GFP tagged PLIN3 fragments was ∼1mg/L. Protein was analyzed by SDS-PAGE, concentrated using a centrifugal tube (10K MWCO, Pall Corporation), aliquoted and stored at -80°C.

### Lipids

The following lipids were purchased from Avanti Polar Lipids: 1,2-dioleoyl-sn-glycero-3- phosphocholine (DOPC, catalog #850375), 1,2-dioleoyl-sn-glycero-3-phosphoethanolamine (DOPE, catalog #850725), 1,2-dioleoyl-sn-glycero-3-phosphate (DOPA, catalog #840875), 1,2- dioleoyl-sn-glycero-3-phospho-L-serine (DOPS, catalog #840035), 1-palmitoyl-2-oleoyl-glycero- 3-phosphocholine (POPC, catalog #850457), 1-palmitoyl-2-oleoyl-sn-glycero-3- phosphoethanolamine (POPE, catalog #850757), 1-palmitoyl-2-oleoyl-sn-glycero-3-phosphate (POPA, catalog #840857), 1-palmitoyl-2-oleoyl-sn-glycero-3-phospho-L-serine (POPS, catalog #840034), 1,2-diphytanoyl-sn-glycero-3-phosphocholine (4ME-PC, catalog #850356), 1,2-diphytanoyl-sn-glycero-3-phosphoethanolamine (4ME-PE, catalog #850402), 1,2-diphytanoyl-sn-glycero-3-phosphate (4ME-PA, catalog #850406), 1,2-diphytanoyl-sn-glycero-3-phospho-L-serine (4ME-PS, catalog #850408), 1-palmitoyl-2-oleoyl-sn-glycerol (DAG, catalog #800815), L- α-phosphatidylinositol (Soy PI, catalog #84044), L-α-phosphatidylinositol-4-phosphate (Brain, Porcine (PI(4)P, catalog #840045), N-stearoyl-D-erythro-sphingosine (C18:1 ceramide, catalog #860518). 1,2,3-Trioleoyl Glycerol was purchase from Cayman Chemicals (TAG, catalog #26950).

### Liposome co-sedimentation assay

To prepare liposomes, lipids were dried under a nitrogen gas stream and dissolved with buffer containing 50mM NaCl and 50mM HEPES pH 8.0 with a total lipid concentration of 2mM, unless otherwise noted. Liposomes were generated by repeated cycles of flash freezing in liquid nitrogen, thawing in 37°C water bath, and vortexing [84]. Liposomes were characterized using dynamic light scattering and showed uniform distribution for most lipid compositions. Lamellarity analysis of liposomes was not carried out, but liposomes likely contained a mixture of multi-lamellar vesicles and unilamellar vesicles. For liposome co-sedimentation assays, 20 μL of 12μM purified protein was incubated with 40μL of 2 mM liposomes for 40mins at 23°C. Protein fractions bound to liposomes were isolated by centrifugation at 100,000 x g for 70mins at 4°C and analyzed by SDS-PAGE.

### Circular Dichroism

Circular dichroism (CD) spectra of purified PLIN3 full length and its constructs were measured on Spectropolarimeter (Jasco, J-715). 0.12∼0.24mg/ml of protein was incubated with 1∼2mM liposomes containing 4ME-PC/PE/PA for 40mins prior to measurement. Liposomes/protein mixture was prepared in the buffer containing 20mM Tris pH 7.5 and 150mM NaCl. CD spectra was measured between 190nm and 260nm in increments of 1nm, with a bandwidth of 50nm and an averaging time of 1min at 25°C. 10 iterations of spectra were averaged and was reported into a mDeg, which was converted to molar ellipticity (m°*M/(10*L*C) of which unit is expressed in degree cm^2^ dmol^-1^. Molar ellipticity of each protein fragments in the absence or presence were plotted using GraphPad prism software. Comparisons of spectra in the presence and the absence of liposome were assessed using unpaired *t* test with *p*<0.05 of statistical significance.

### Artificial lipid droplet flotation assay

Artificial lipid droplets (ALDs) were prepared by mixing phospholipids and triacylglycerols (TAGs). Di-oleoyl phospholipids were dried under a nitrogen gas stream. Dried lipids were resuspended with lipid droplet flotation assay buffer containing 50mM NaCl and 50mM HEPES pH 8.0 and TAGs in the molar ratio of 2:5. ALDs were formed by repeating the cycles of vortex for 10sec and rest for 10sec. ALDs were further vortexed before use for the assay. 100μL of ALDs were mixed with 5μL of human PLIN3 and 105μL of lipid flotation assay buffer to give a final concentration of 5μM protein and 1.5mM phospholipids. The average diameters of different ALDs were measured by dynamic light scattering (DLS). The concentration of ALDs were determined by optical density at 600nm. The mixture of human PLIN3 and ALDs was incubated for 1h at 23°C. To generate a sucrose gradient, 140μL of a 75% sucrose solution was added to 210μL of the protein-ALD mixture to give a final concentration of 30% sucrose. The resulting mixture (320μL) was transferred to an ultra-centrifuge tube. 260μL of a 25% sucrose solution was applied on top of 30% sucrose/protein/ALDs mixture. Lastly, 60μL of lipid flotation assay buffer was laid on the top. ALDs were floated by centrifugation at 76,000 x g for 3h at 20°C with sucrose gradient of 30% (bottom), 25% (middle), and 0% (top). Three fractions of 100μL from the top, 260μL from the middle and 280μL from the bottom were collected and analyzed by SDS-PAGE.

### Hydrogen-Deuterium Exchange Mass Spectrometry (HDX-MS)

#### Sample preparation

HDX reactions for PLIN3 deuterium pulse were conducted in a final reaction volume of 10μL with a final concentration 2.12μM PLIN3. Exchange was carried out in triplicate for a single time point (3 sec at 0 °C). Hydrogen deuterium exchange was initiated by the addition of 9μL of D2O buffer solution (20mM HEPES pH 7.5, 100mM NaCl) to 1μL of protein, to give a final concentration of 84.9% D2O. Exchange was terminated by the addition of 60μL acidic quench buffer at a final concentration 0.6M guanidine-HCl and 0.9% formic acid. Samples were immediately frozen in liquid nitrogen at -80°C. Fully deuterated samples were generated by first denaturing the protein in 3M guanidine for 1h at 20°C. Following denaturing, 9μL of D2O buffer was added to the 1μL of denatured protein and allowed to incubate for 10mins at 20°C before quenching with 0.6M guanidine-HCl and 0.9% formic acid. Samples were immediately frozen in liquid nitrogen at -80°C, and all timepoints were created and run in triplicate.

HDX reactions comparing PLIN3 in the presence of PA liposomes (60mol% 4ME-PC, 20mol% 4ME-PE, 20mol% 4ME-PA) were conducted in a final reaction volume of 10μL with a final protein concentration of 3μM and final liposome concentration of 1mM. Protein and liposomes were preincubated together for 2mins at 20°C before the addition of 7μL D2O buffer solution (50mM HEPES pH 8.0, 50mM NaCl) for a final concentration of 63% D2O. Exchange was carried out for 3sec, 30sec, 300sec and 3000sec, and exchange was terminated by the addition of 60μL acidic quench buffer at a final concentration 0.6M guanidine-HCl and 0.9% formic acid. Samples were immediately frozen in liquid nitrogen at -80°C, and all timepoints were created and run in triplicate.

HDX reactions comparing PLIN3 in the presence of two different liposomes (60mol% DOPC, 20mol% DOPE, 20mol% DAG liposomes and 80mol% 4ME-PC, 20mol% 4ME-PE liposomes) were conducted in a final reaction volume of 10μL with a final protein concentration of 3μM and final liposome concentration of 500μM. Protein and liposomes were preincubated together for 2mins at 20°C before the addition of 8μL D2O buffer solution (50mM HEPES pH 8.0, 50mM NaCl) for a final concentration of 72% D2O. Exchange was carried out for 3sec, 30sec, 300sec and 3000sec, and exchange was terminated by the addition of 60μL acidic quench buffer at a final concentration 0.6M guanidine-HCl and 0.9% formic acid. Samples were immediately frozen in liquid nitrogen at -80°C, and all timepoints were created and run in triplicate.

#### Protein digestion and MS/MS data collection

Protein samples were rapidly thawed and injected onto an integrated fluidics system containing a HDx-3 PAL liquid handling robot and climate-controlled (2°C) chromatography system (LEAP Technologies), a Dionex Ultimate 3000 UHPLC system, as well as an Impact HD QTOF Mass spectrometer (Bruker). The protein was run over either one (at 10°C) or two (at 10°C and 2°C) immobilized pepsin columns (Trajan; ProDx protease column, 2.1mm x 30mm PDX.PP01-F32) at 200μL/min for 3mins. The resulting peptides were collected and desalted on a C18 trap column (Acquity UPLC BEH C18 1.7mm column (2.1 x 5mm); Waters 186003975). The trap was subsequently eluted in line with an ACQUITY 1.7μm particle, 100 × 1mm2 C18 UPLC column (Waters), using a gradient of 3-35% B (Buffer A 0.1% formic acid; Buffer B 100% acetonitrile) over 11mins immediately followed by a gradient of 35-80% over 5mins. Mass spectrometry experiments acquired over a mass range from 150 to 2200m/z using an electrospray ionization source operated at a temperature of 200°C and a spray voltage of 4.5kV.

#### Peptide identification

Peptides were identified from the non-deuterated samples of PLIN3 using data-dependent acquisition following tandem MS/MS experiments (0.5sec precursor scan from 150-2000m/z; twelve 0.25sec fragment scans from 150-2000m/z). MS/MS datasets were analyzed using PEAKS7 (PEAKS), and peptide identification was carried out by using a false discovery based approach, with a threshold set to 1% using a database of known contaminants found in Sf9 and E. coli cells [85]. The search parameters were set with a precursor tolerance of 20ppm, fragment mass error 0.02 Da, charge states from 1-8, leading to a selection criterion of peptides that had - 10logP scores of 22.8 for the pulse experiment and 20.9 for the liposome experiment.

#### Mass Analysis of Peptide Centroids and Measurement of Deuterium Incorporation

HD-Examiner Software (Sierra Analytics) was used to automatically calculate the level of deuterium incorporation into each peptide. All peptides were manually inspected for correct charge state, correct retention time, appropriate selection of isotopic distribution, etc. Deuteration levels were calculated using the centroid of the experimental isotope clusters. Results are presented as relative levels of deuterium incorporation, with the only correction being applied correcting for the deuterium oxide percentage of the buffer utilized in the exchange (63% and 72%). For the experiment with a fully deuterated sample, corrections for back exchange were made by dividing the pulse %D value by the fully deuterated %D value and multiplying by 100. The raw %D incorporation for the fully deuterated sample is included in the source data, with the average back exchange being 33%, and ranging from 10-60%. Differences in exchange in a peptide were considered significant if they met all three of the following criteria: : ≥5% change in exchange and ≥0.4 Da difference in exchange. The raw HDX data are shown in two different formats. Samples were only compared within a single experiment and were never compared to experiments completed at a different time with a different final D2O level. The data analysis statistics for all HDX-MS experiments are in Supplemental Table 1 according to the guidelines of [86]. The mass spectrometry proteomics data have been deposited to the ProteomeXchange Consortium via the PRIDE partner repository with the dataset identifier PXD025717 [87].

#### Dynamic light scattering

The size of liposomes was measured by dynamic light scattering (Brookhaven Instruments NanoBrook Omni), with or without full length PLIN3. Liposomes were prepared in the buffer containing 50mM HEPES pH 8.0 and 50mM NaCl by repeating the freeze/thaw cycles to prevent multilamellar vesicle formation. Each protein construct was incubated with liposomes in 1:200 of molar ratio at 25°C for 40mins. DLS was conducted using a scattering angle of 90°, and the liposome/protein mixture was equilibrated for 5mins after loading into the instrument to get a uniform temperature and minimize any loading effects prior to measurement. The number average size distribution (%) was considered as a relative concentration of particles with a certain size. For analysis, measurements with large outlier peaks, which were suspected to be due to the dust particles or aggregated vesicles, were discarded, and three runs that did not contain outlier peaks were used for the data analysis. The mode of the distribution (e.g., the size having the highest peak in the number average size distribution) was chosen from each of these three measurements and averaged to obtain the reported average diameter. A representative size distribution is shown for each sample.

#### Site-directed spin-labeling

Recombinant PLIN3 cysteine-substituted proteins were purified using Ni-NTA resin in a buffer 500mM NaCl, 20mM Tris pH 7.5, 5% glycerol and eluted in a buffer of 500mM NaCl, 20mM Tris pH 7.5, 5% Glycerol, 300mM imidazole. Elution was spin-labeled with 1μg/mL (S-(1-oxyl-2,2,5,5- tetramethyl-2,5-dihydro-1H-pyrrol-3-yl) methyl methanesulfonothioate), MTSL, (Santa Cruz Biotech) dissolved in acetonitrile at 4^°^C overnight. The spin-labeled proteins were further purified by HiLoad 26/600 Superdex 200 pg column (GE life sciences) in a buffer of 150mM NaCl, 20mM Tris pH 7.5 to remove unreacted spin labels.

#### Sample preparations for DEER spectroscopy

Protein samples in solutions were prepared at ca. 40-60μM concentrations in 20mM Tris at pH 7.5, 150mM NaCl, and 40% glycerol. Liposome samples containing 10μM protein and 3.33 mM lipid were prepared by mixing 30μM protein stocks with 5mM liposome solutions using 1 to 2 aliquots followed by 30mins incubation at room temperature. The liposomes were freshly prepared by rehydration of lyophilized 80/20 mol% 4ME-PC/PE followed by seven freeze/thaw cycles. Freshly-prepared 20μL of spin-labeled samples were loaded into 1.8mm inner diameter Pyrex sample tubes (Wilmad-LabGlass), frozen by plunging in liquid nitrogen, and stored in liquid nitrogen for PDS measurements.

#### DEER data collection and analysis

PDS DEER measurements were performed at 60K with a home-built Ku band pulse ESR spectrometer operating around 17.3GHz [64, 88]. A four-pulse DEER sequence [89] was employed using for spin-echo detection *π*/2- and *π*-pulses with respective widths of 16 and 32ns, the *π*-pulse for pumping was 16ns. The detection frequency matched the peak at the low-field spectral edge, while pumping was performed at a lower by 70MHz frequency, corresponding to the central maximum. A 32-step phase cycle [90] was applied to suppress unwanted contributions to the signal. Nuclear electron spin-echo envelope modulation (ESEEM) caused by surrounding protons was suppressed using the data from four measurements. In this method the initial pulse separations and the start of detection were advanced by 9.5ns in subsequent measurements, i.e. by the quarter period of the 26.2MHz nuclear Zeeman frequency for protons at 0.615T corresponding to the working frequency. The four signal traces were summed to achieve deep suppression of nuclear ESEEM, [91].

The solution samples had phase memory times, *T*m*’*s, of about 2.5μs, so the data could be recorded up to 3μs evolution time (*t*m). Such evolution times did not provide sufficient decay of the dipolar coherence to the background level, providing however clear indication of unstructured nature of the protein in solution. Any residual secondary structure could be probed by using multiple labeling sites, however other techniques did not encourage this undertaking. For solutions the DEER data were acquired in less than 12h. Faster phase relaxation times (∼1μs) and low spin concentrations in liposome samples required signal averaging for ∼24h to facilitate good reconstruction by Tikhonov regularization. [92, 93] For liposomes, time domain DEER data, *V(t*), were recorded and preprocessed using standard approaches [62, 65, 89, 94] before their reconstruction into distance distributions, *P*(*r*)’s. The first step is to remove signal decay caused by intermolecular dipole-dipole couplings, followed by subtracting the residual background from the spins whose partner was not flipped by the pump or missing. This was done, as usual, by fitting the latter points (about a half of the record) of ln[*V(t*)] to a low-order polynomial, usually a straight line for solutions (and often for liposomes), extrapolating it to zero time, and subtracting out from ln[*V(t*)]; so that the antilog yields *u*(*t*) = *d(t*) + 1. Here *d*(*t*) is the dipolar evolution representing part. Once *u*(*t*) is normalized as *u*(*t*)/*u(*0), it serves as a typical form of DEER data presentation, while *v*(*t*)=(*u*(*t*)-1)/*u*(0), gives background-free “dipolar” data to be converted to a distance distribution between spins in pairs. We used L-curve Tikhonov regularization [92] for distance reconstruction. The Tikhonov regularization utility allowed selection of either the signal or its derivatives in the Tikhonov functional and selection of the regularization parameter. The latter increases *P*(*r*) smoothness at the expense of introducing some broadening.

Dipolar signal amplitude (“modulation depth”) is given by *v*(0). This would be accurate to the extent the background or *V*(∞), the asymptotic value of *V(t*), is known. This depth, tabulated by calculations for typical pulse sequence setup and verified in numerous experiments, is a measure of a fraction of spins in pairs or oligomers [69, 95, 96]. It is thus useful in estimate of spin-labeling efficiency and protein concentration.

#### Cell culture

Cos7 cells were maintained in Dulbecco’s modified Eagle’s medium (DMEM) supplemented with 10% heat inactivated fetal bovine serum (Life Technologies), 4.5gL^-1^ D-glucose, 0.1gL^-1^ sodium pyruvate (Life Technologies) and 1% penicillin-streptomycin (Life Technologies). The cells were cultured at 37°C under a 5% CO2 atmosphere.

#### Transfection

When indicated, Cos7 cells (60–70% confluence) plated into a 35mm cell-culture Mattek dishes (with a glass coverslip at the bottom), (MatTek Corp. Ashland, MA). were transfected with 3μg of plasmid DNA/ml using Polyethylenimine HCl MAX (Polysciences) following the manufacturer’s instructions. For co-expressions, RFP-KDEL or GFP-Plin3 constructs in equal concentrations (1.5–2μg for each one) were transfected to cells 24h prior observation.

#### Giant unilamellar vesicles, GUVs

GUVs were prepared by electroformation following the protocol described in [97]. Phospholipids, dioleoyl phosphatidylcholine (PC) (70%) and dioleoyl phosphatidylethanolamine (PE) (29%), Rhodamine-DOPE 1% (w/w)), were purchased from Avanti Polar Lipids. The lipid mixture, in chloroform at 0.5mM, was dried on an indium tin oxide (ITO)-coated glass plate. The lipid film was desiccated for 1h. The chamber was sealed with another ITO- coated glass plate. The lipids were then rehydrated with a sucrose solution (275mOsm). Electroformation was done using 100Hz AC voltage at 1.0 to 1.4Vpp and maintained for at least 1h.

#### Droplet-embedded vesicles (DEVs) preparation

Artificial LDs (aLDs) were prepared in HKM buffer: 50mM HEPES, 120mM Kacetate, and 1mM MgCl2 (in Milli-Q water) at pH 7.4 and 275±15mOsm. To do so, 5μL of the lipid oil solution (triolein or diacylglycerol purchased from Sigma) was added to 45mL of HKM buffer and the mixture was sonicated as described in [98]. The resulting emulsion was then mixed with GUVs for five minutes under rotator to generate DEVs [98]. DEVs were then placed on a glass coverslip pretreated with 10%(w/w) BSA and washed three times with buffer.

#### Protein binding to DEVs

For the protein binding experiments, 50μL HKM were deposited on the BSA-treated glass, 30μL of the DEV solution added and 1.5μL purified PLIN3 or PLIN3 fragments added (in the case of 107-204 and 1-106 fragments, 3μL were added). The final protein concentration in the solution was between 1μM to 4μM depending on the protein. For quantifications, 10-15 DEVs were considered and the signal on the droplet and bilayer was determined by Fiji, via drawing 5-10 points-thick line profiles.

#### Neutral lipid synthesis induction

Wherever relevant, cells were exposed for 1h to 350μM oleic acid (OA) coupled to bovine serum albumin (BSA) (0.2% weight/volume) to induce neutral lipids’ synthesis. LipidTox DeepRed (Thermo Fischer), was used to visualize lipid droplets or membranes enriched in neutral lipids.

#### Giant Intra-Cellular ER vesicle experiments

For GERV experiments, Cos7 cells were first transfected for 24h with the indicated eGFP plasmids and RFP-KDEL. The culture media of the cells was next replaced by a hypotonic culture media (DMEM:H2O, 1:20). The cells were then incubated at 37°C, 5% CO2 for 5mins, to induce GERVs, and then they were visualized. Neutral lipids’ synthesis was triggered by the addition of OA and Z-stacks imaging of different entire cells was performed before and 15mins after OA administration, as described in [77]. For the DGAT1 (Sigma, PF-04620110) and DGAT2 (Sigma, PF-06424439) inhibitors used in Cos7, the dilution applied was 1/1000 for a final concentration of 3μg/mL. In the GERV protocol, the inhibitors were added to the cell medium before cell transfection or during the hypotonic solution addition. The signal of PLIN3 or fragments thereof was quantified for tens of GERVs by depicting the mean signal on the GERVs to which was subtracted the bulk cytosolic signal; this resulted signal was normalized to the cytosolic signal.

#### Structure prediction by AlphaFold and RoseTTAFold

Structure prediction of PLIN3 was carried out by AlphaFold and RoseTTAFold. Two different confidence values were generated (pLDDT from AlphaFold and rmsd from RoseTTAFold). These values were converted using PHENIX to get relative confidence level from two softwares, AlphaFold and RoseTTAFold [99].

## Acknowledgements

This work was supported by in part by the NIGMS (R35GM128666, MVA), the Sloan Research Foundation (MVA), the Natural Science and Engineering Research Council of Canada (Discovery Grant 2020-04241, JEB) with salary support from the Michael Smith Foundation for Health Research (Scholar award 17868, JEB), the ANR (18-CE11-0012-01-MOBIL, CE11-0032- 02-LIPRODYN and 21-CE13-0014-LIPDROPER, ART), and the NSF (CBET 1903189 and DGE 1922639, SRB). ESR study, conducted at ACERT, was supported by NIH/NIGMS grants 1R24GM146107 and P41GM103521.

## Author contributions

YMC generated constructs for E. coli and mammalian cell expression, purified all proteins, conducted liposome sedimentation, artificial lipid droplet binding assays, CD, DLS with assistance from SF, prepared figures, and wrote the initial draft. DA conducted all DEV and cell culture experiments. KDF, MLJ, and BEM setup, analyzed, and/or prepared figures and text from the HDX-MS data. PPB designed, acquired, analyzed, and prepared figures for all PDS- ESR data. JEB, ART, JHF, SRB and MVA supervised the work and provided funding support.

---

**Supplementary Figure 1.**
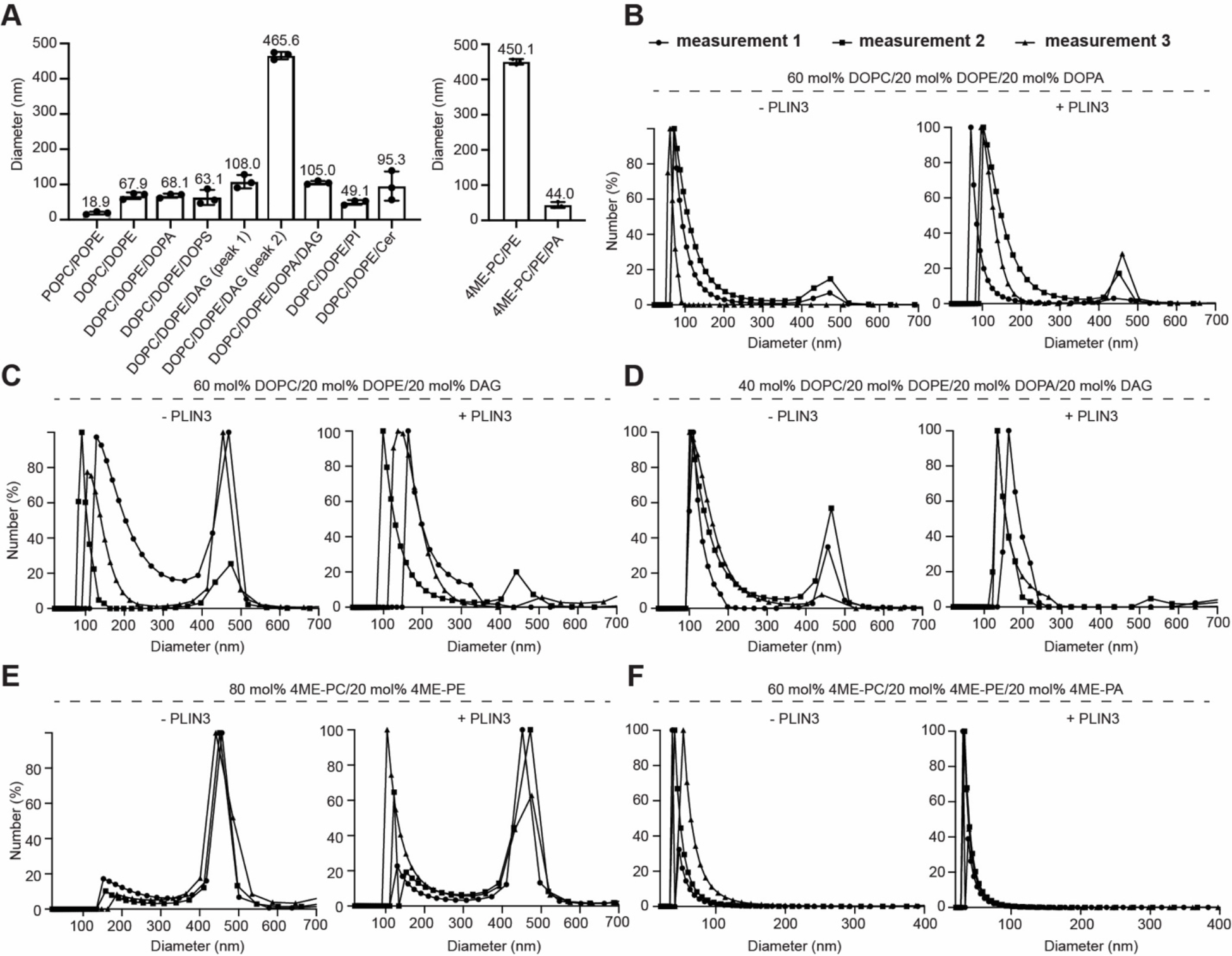
**A)** DLS analysis of liposomes with different lipid composition. For each liposome, three measurements were plotted. **B-F)** DLS analysis of different liposomes with or without full length PLIN3. For each liposome, the number average size distribution (%) was recorded and plotted (n=3) in the presence and absence of full length PLIN3.

**Supplementary Figure 2.**
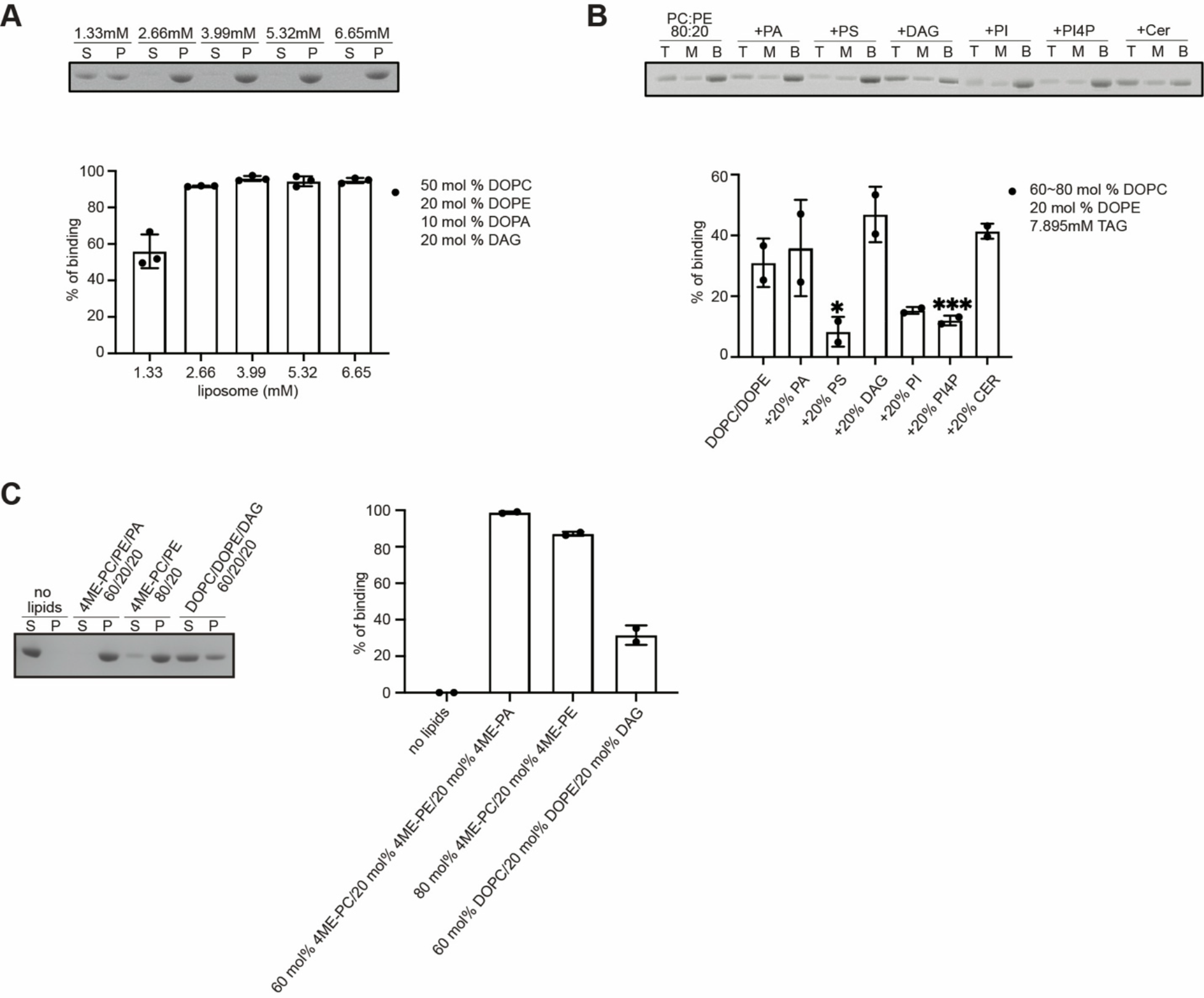
**A)** SDS-PAGE and quantitative analysis of human PLIN3 recruitment by increasing the total amount of liposomes. The molar ratio of DOPC/DOPE/DOPA/DAG in 50:20:10:20 was kept same. Lane S represents unbound human PLIN3 from supernatant. Lane P represents pelleted human PLIN3 that bound to liposomes. Bar graph shows total increase in liposome amount results almost 100% recruitment of PLIN3 to liposome. **B)** SDS-PAGE and quantitative analysis of human PLIN3 recruitment to ALDs generated with DO-phospholipids and TAGs (C16) by adding additional with 20 mol% of lipids such as PS, PA, DAG, PI, PI4P and ceramide. Top, middle and bottom fractions after sucrose-gradient centrifuge were indicated as T, M and B, respectively. Statistical analysis was performed using ordinary one-way ANOVA with Tukey’s multiple comparison test (n=2, *, p=0.0258, ***,p=0.004). **C)** SDS-PAGE and quantitative analysis of human PLIN3 recruitment to liposomes in buffer containing 100mM NaCl and 20mM HEPES pH 7.0. Three different liposomes that were applied in HDX-MS were generated. Lane S represents unbound human PLIN3 from supernatant. Lane P represents pelleted human PLIN3 that bound to liposomes.

**Supplementary Figure 3.**
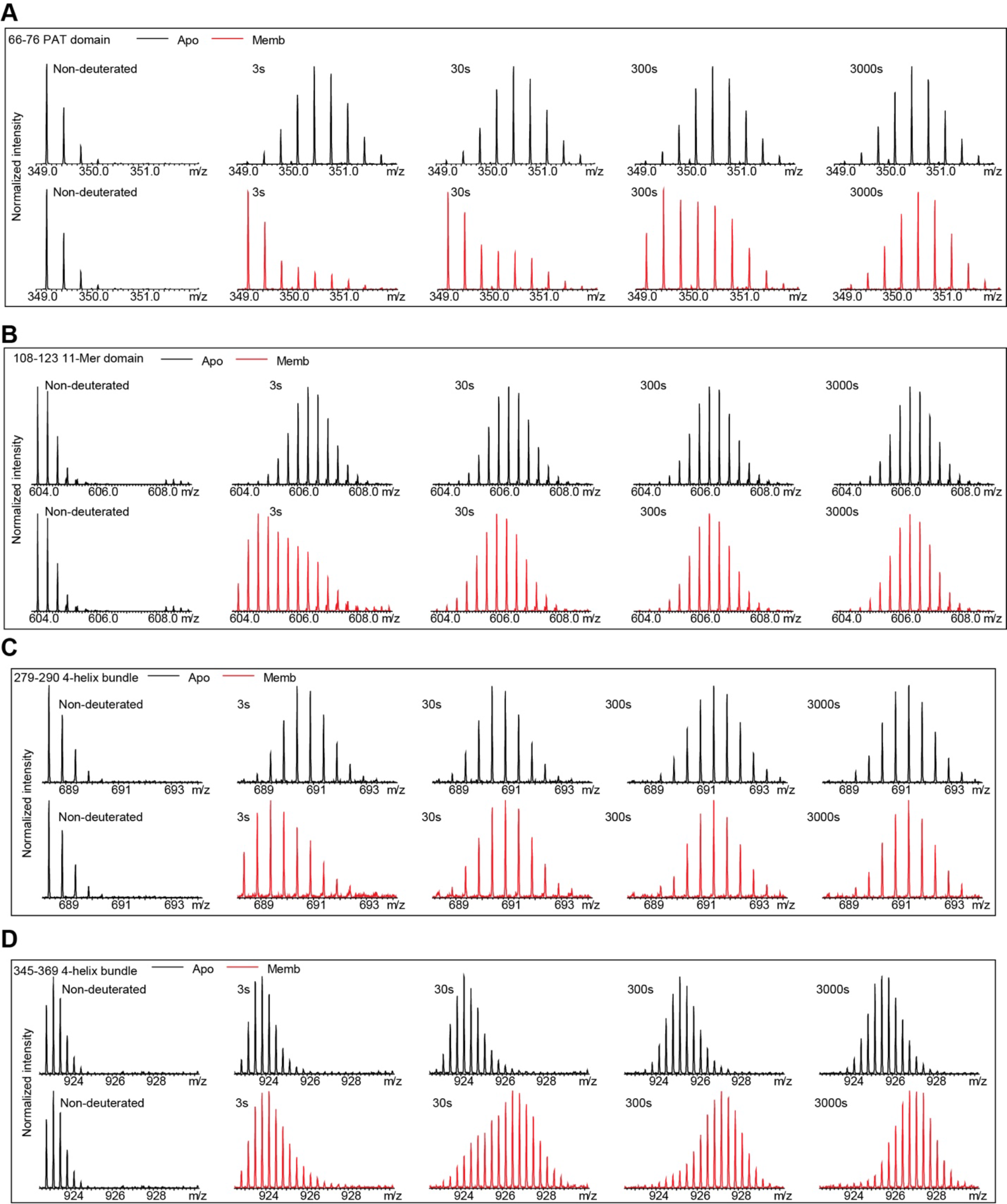
Representative bimodal distribution mass spectra from the peptides at PAT domain, 11-mer repeats, and 4-helix bundle of PLIN3 after deuterium exchange.

**Supplementary Figure 4.**
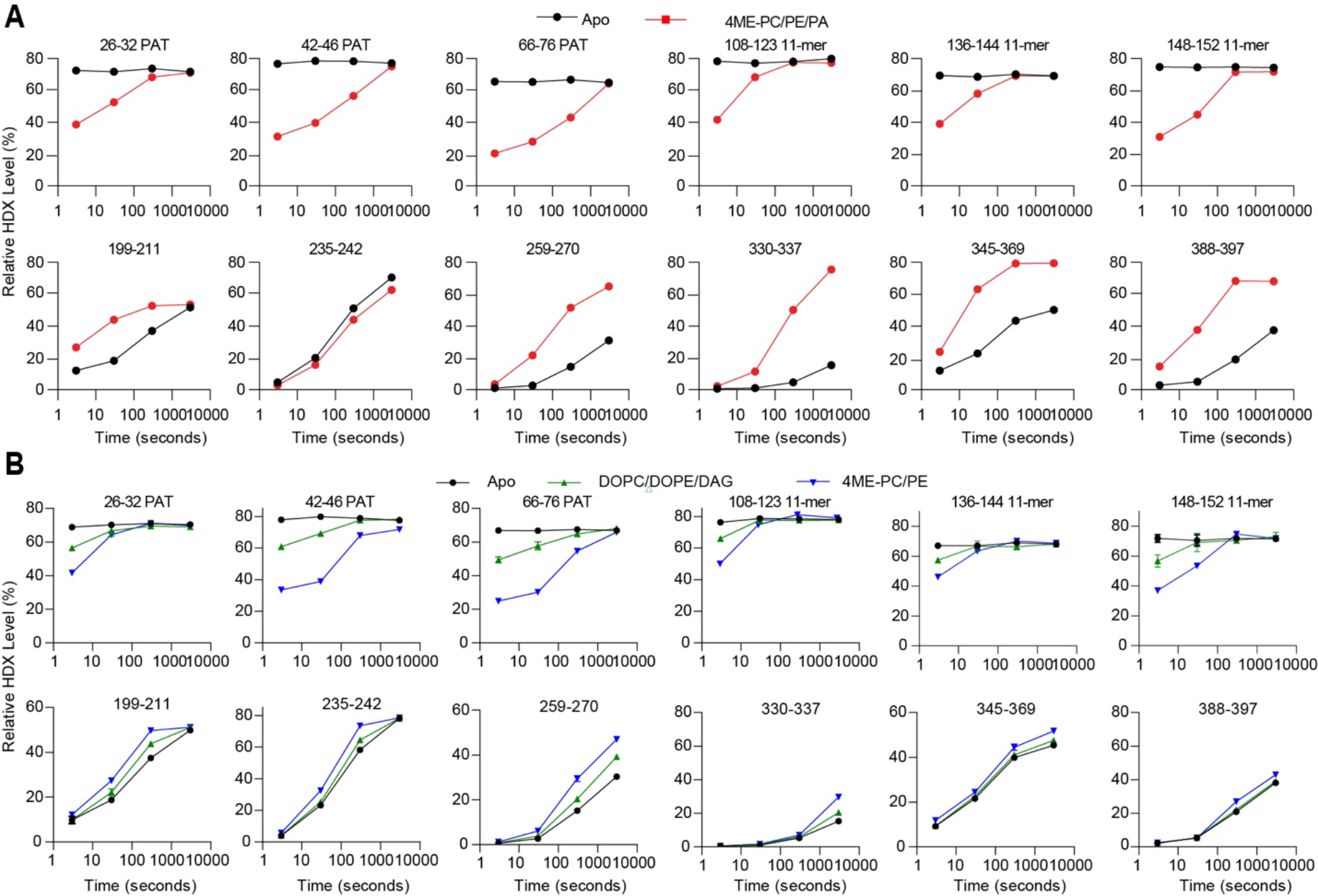
**A)** Deuterium exchange uptake plots for selected peptides in the PAT domain, 11-mer repeats and 4-helix bundle were plotted across all timepoints as 3s, 30s, 300s and 3000s. Data points in the absence and presence of liposomes were colored in black and red, respectively. Liposomes were generated with 60 mol% 4ME-PC, 20 mol% 4ME-PE and 20 mol% 4ME-PA). All peptides are shown in the source data. **B)** Deuterium exchange uptake plots for selected peptides in the presence of two different liposomes are plotted across all timepoints as 3s, 30s, 300s and 3000s colored according the the legend. All peptides are shown in the source data.

**Supplementary Figure 5.**
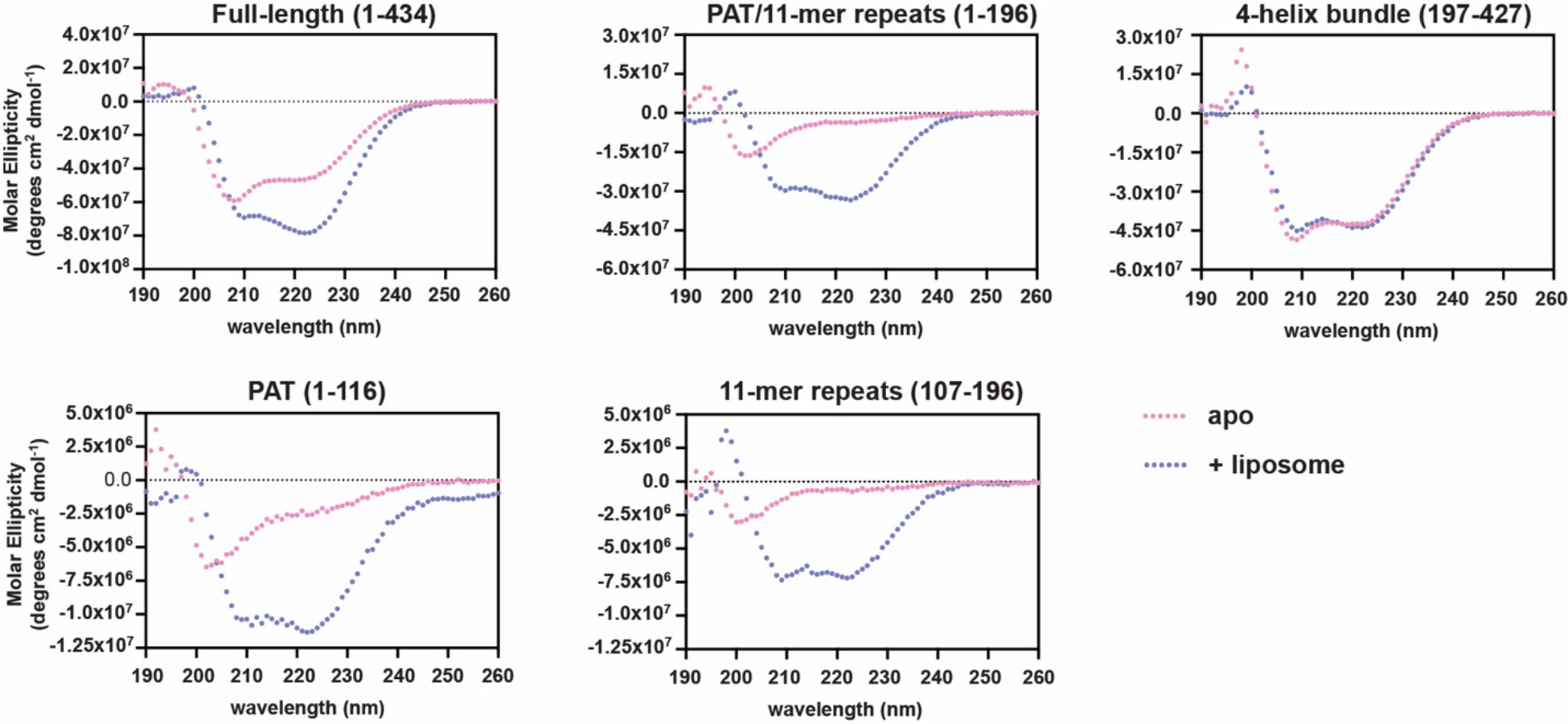
CD analysis of PLIN3 full length and various fragments with or without 4ME-PC/PE/PA liposomes

**Supplementary Figure 6.**
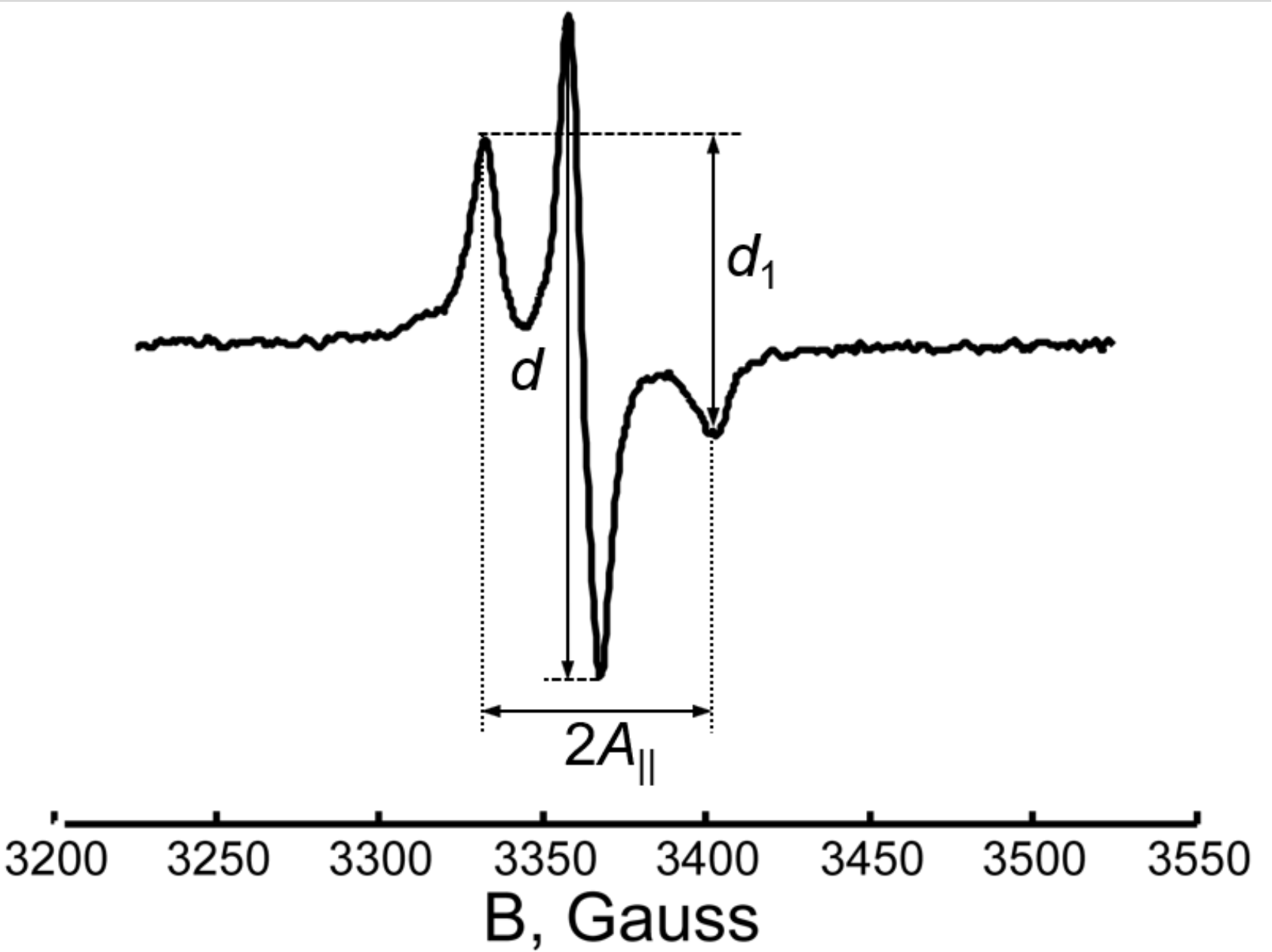
CW ESR spectrum of 37C/144C in lipid recorded at 9,434 GHz frequency. The Parameter *Δ*, defined [100] as the ratio of d1/d, is 0.45, which for the MTSL spin label indicates the distance in range of 1.5-2.0 nm [100, 101], i.e. somewhat shorter than reported by DEER which has reduced sensitivity to distances below 2.0 nm.

**Supplementary Figure 7.**
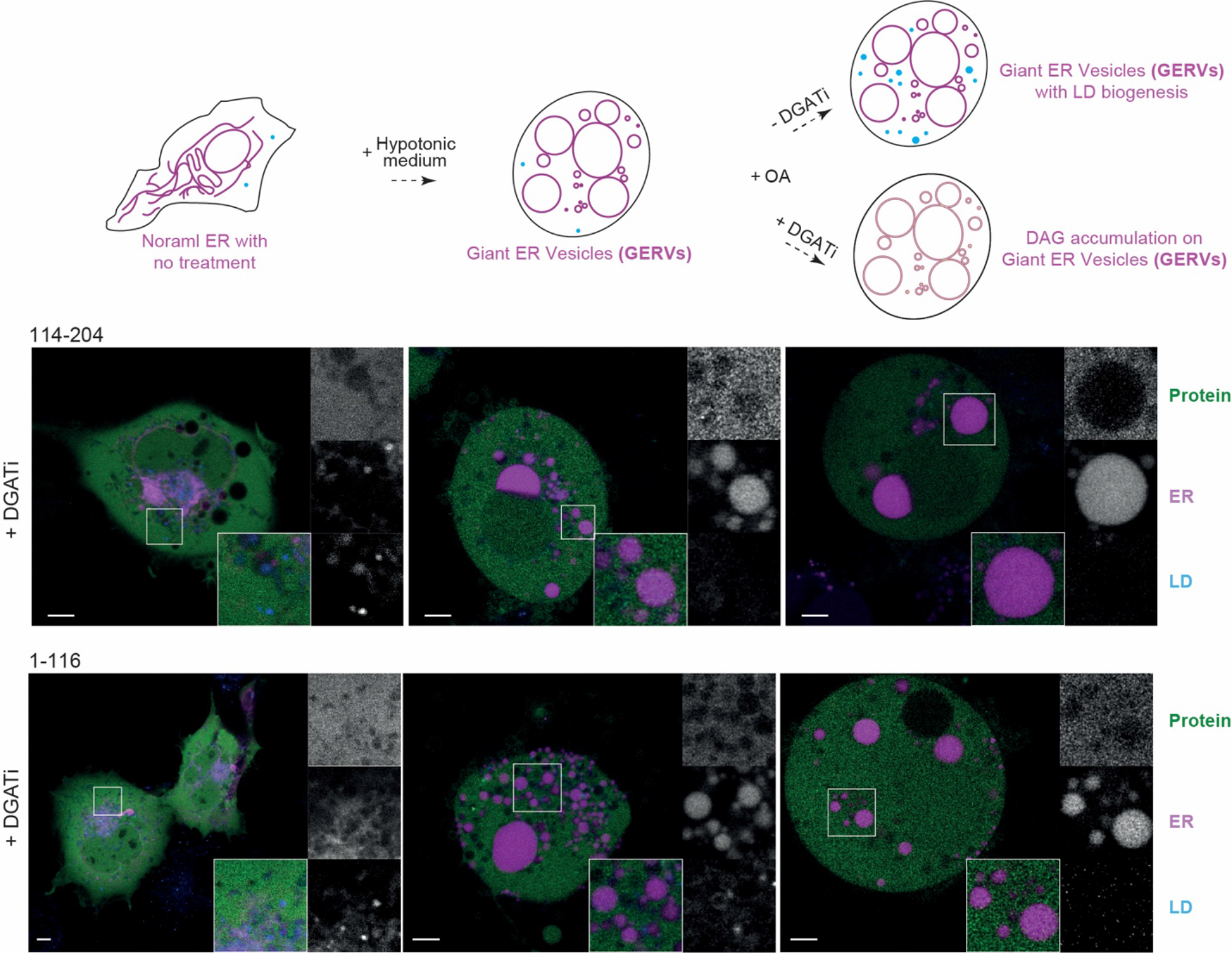
Subcellular localization of GFP tagged 114-204 and 1-116 constructs of PLIN3 was visualized in green in Cos7 cells under fluorescent microscope ZEISS LSM800 Airyscan. ER was visualized with ER specific marker RFP-KDEL in magenta. Lipid droplets were labeled with LipidTox Deepred to stain neutral lipids in cyan. After treated with hypotonic medium, cells were supplemented with oleate in the presence or absence of DGAT1/2 inhibitors.

**Supplementary Table 1.** Summary of all HDX-MS data processing.

|  |  |  |  |  |  |  |
| --- | --- | --- | --- | --- | --- | --- |
| Data set | PLIN3 order disorder Expt-1 | PLIN3 Apo Expt-2 | PLIN3 + 60% 4ME-PC, 20% 4ME-PE, 20% 4ME-PA liposomes Expt-2 | PLIN3 Apo Expt-3 | PLIN3 + 60% DOPC, 20% DOPE, 20% DAG liposomes Expt-3 | PLIN3+ + 80% 4ME-PC, 20% 4ME-PE liposomes Expt-3 |
| HDX reaction details | %D2O=84.9 % pH(read)=7.5 Temp=20°C | %D2O=63% pH(read)=8.0 Temp=20°C | %D2O=63% pH(read)=8.0 Temp=20°C | %D2O=72% pH(read)=8.0 Temp=20°C | %D2O=72% pH(read)=8.0 Temp=20°C | %D2O=72% pH(read)=8.0 Temp=20°C |
| HDX time course (seconds) | 0.3s, fully deuterated | 3s, 30s, 300s, 3000s | 3s, 30s, 300s, 3000s | 3s, 30s, 300s, 3000s | 3s, 30s, 300s, 3000s | 3s, 30s, 300s, 3000s |
| HDX controls | FD | N/A | N/A | N/A | N/A | N/A |
| Back-exchange | Corrected by fully deuterated sample | No correction, deuterium levels are relative | No correction, deuterium levels are relative | No correction, deuterium levels are relative | No correction, deuterium levels are relative | No correction, deuterium levels are relative |
| Number of peptides | 144 | 90 | 90 | 90 | 90 | 90 |
| Sequence coverage | 99.8% | 91.7% | 91.7% | 91.7% | 91.7% | 91.7% |
| Average peptide /redundancy | Length= 17.3 Redundancy = 5.4 | Length= 13.0 Redundancy = 2.6 | Length= 13.0 Redundancy = 2.6 | Length= 13.0 Redundancy = 2.6 | Length= 13.0 Redundancy = 2.6 | Length= 13.0 Redundancy= 2.6 |
| Replicates | 3 | 3 | 3 | 3 | 3 | 3 |
| Repeatability | Average StDev=0.4% | Average StDev=0.6% | Average StDev=0.6% | Average StDev=0.8% | Average StDev=1.0% | Average StDev=0.9% |
| Significant differences in HDX | N/A | >5% and >0.4 Da and unpaired t-test $\leq 0.01$ | >5% and >0.4 Da and unpaired t-test $\leq 0.01$ | >5% and >0.4 Da and unpaired t-test $\leq 0.01$ | >5% and >0.4 Da and unpaired t-test $\leq 0.01$ | >5% and >0.4 Da and unpaired t-test $\leq 0.01$ |

## Supplementary Information 1

Codon optimized DNA sequence of human perilipin 3

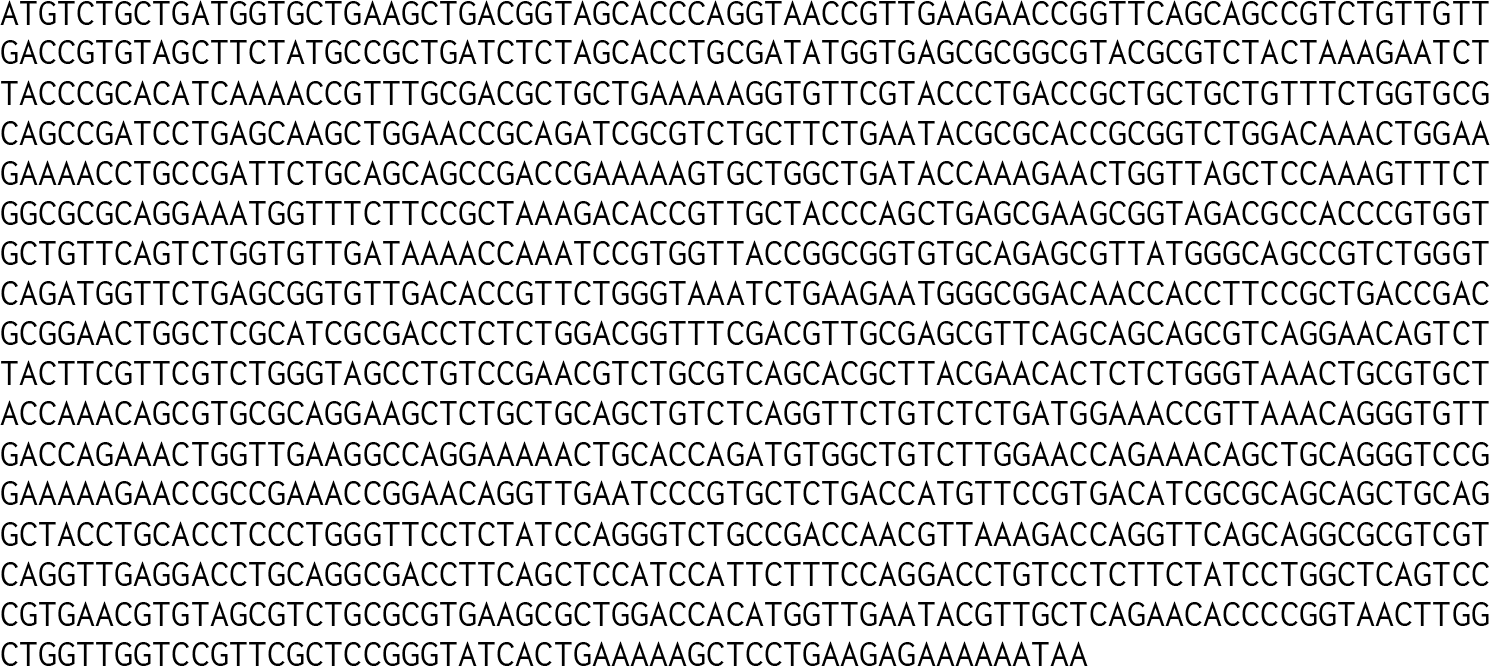

